# The ABCF ATPase New1 resolves translation termination defects associated with specific tRNA^Arg^ and tRNA^Lys^ isoacceptors in the P site

**DOI:** 10.1101/2024.05.29.596377

**Authors:** Kathryn Turnbull, Helge Paternoga, Esther von der Weth, Artyom A. Egorov, Agnieszka A. Pochopien, Yujie Zhang, Lilit Nersisyan, Tõnu Margus, Marcus J.O. Johansson, Vicent Pelechano, Daniel N. Wilson, Vasili Hauryliuk

## Abstract

The efficiency of translation termination is determined by the nature of the stop codon as well as its context. In eukaryotes, recognition of the A-site stop codon and release of the polypeptide are mediated by release factors eRF1 and eRF3, respectively. Translation termination is modulated by other factors which either directly interact with release factors or bind to the E-site and modulate the activity of the peptidyl transferase center. Previous studies suggested that the *Saccharomyces cerevisiae* ABCF ATPase New1 is involved in translation termination and/or ribosome recycling, however, the exact function remained unclear. Here, we have applied 5PSeq, single-particle cryo-EM and readthrough reporter assays to provide insight into the biological function of New1. We show that the lack of New1 results in ribosomal stalling at stop codons preceded by a lysine or arginine codon and that the stalling is not defined by the nature of the C-terminal amino acid but rather by the identity of the tRNA isoacceptor in the P-site. Collectively, our results suggest that translation termination is inefficient when ribosomes have specific tRNA isoacceptors in the P-site and that the recruitment of New1 rescues ribosomes at these problematic termination contexts.

## INTRODUCTION

Translation termination and ribosome recycling are the two last steps of the protein synthesis on the ribosome (Dever & Green, 2012; Hellen, 2018). In eukaryotes, termination is governed by two release factors – eRF1 and eRF3 (Inge-Vechtomov *et al*, 2003; Zhouravleva *et al*, 1995) – which in yeast *Saccharomyces cerevisiae* are encoded by the *SUP45* and *SUP35* genes, respectively (Stansfield *et al*, 1995; Stansfield & Tuite, 1994). Acting as a structural mimic of tRNA, eRF1 recognises the stop codon in the ribosomal A-site and catalyses the release of the polypeptide (Brown *et al*, 2015; Matheisl *et al*, 2015). eRF3 is a translational GTPase (Atkinson, 2015; Atkinson *et al*, 2008) that delivers eRF1 to the ribosome (Alkalaeva *et al*, 2006) and increases the efficiency of eRF1-mediated termination (Eyler *et al*, 2013; Salas-Marco & Bedwell, 2004). Following GTP hydrolysis and the departure of eRF3, ABCE ATPase ribosome recycling factor ABCE1/Rli1 accommodates in the ribosomal GTPase centre and establishes interactions with eRF1 (Brown *et al*., 2015; des Georges *et al*, 2014; Shao *et al*, 2016; Taylor *et al*, 2012). ABCE1/Rli1 then splits the 80S ribosome into a free 60S subunit and a tRNA/mRNA-bound 40S subunit (Khoshnevis *et al*, 2010; Shoemaker & Green, 2011). Finally, the mRNA and deacylated tRNA are disassociated from the 40S subunit by Tma64 (eIF2D) or the Tma20/Tma22 (MCT-1/DENR) complex (Skabkin *et al*, 2010; Young *et al*, 2018).

The identity of the stop codon as well as sequence context surrounding it strongly influence the efficiency of translation termination (Bonetti *et al*, 1995; Tate *et al*, 1996). In eukaryotes, the fidelity of stop codon recognition follows the ranking UAA > UAG > UGA (Bonetti *et al*., 1995; Cridge *et al*, 2018). Termination at weak stop codons within a weak context is more sensitive to perturbations that globally decrease the termination efficiency, such as depletion of eRF1 (Cridge *et al*., 2018). The key stop codon context determinant is the +4 position, i.e. the 3’ nucleotide following the stop codon, which together with the stop codon itself forms the so-called the “extended stop codon” (Anzalone *et al*, 2019; Cridge *et al*., 2018; Harrell *et al*, 2002; Mangkalaphiban *et al*, 2024; Mangkalaphiban *et al*, 2021; McCaughan *et al*, 1995; Namy *et al*, 2001; Tate *et al*, 1995; Wangen & Green, 2020). The UGA stop codon followed by a 3’ cytosine is the least efficient in promoting termination (Anzalone *et al*., 2019; Bonetti *et al*., 1995; Tate *et al*., 1996). Structures of eRF1 locked on ribosomal termination complexes revealed that both the stop codon and the following nucleotide occupy the A-site and are “read” by the release factor, thus explaining the crucial role of the +4 position (Brown *et al*., 2015; des Georges *et al*., 2014; Shao *et al*., 2016; Taylor *et al*., 2012). The nucleotide context 5’ of the stop codon also affects termination efficiency. Experiments with a series of readthrough reporters suggested that in *S. cerevisiae* the adenines in positions -1 and -2, such as in CAA (encoding glutamine) and GAA (encoding glutamic acid), could serve as determinant of high stop codon readthrough, i.e. low fidelity of stop codon recognition (Tork *et al*, 2004). However, a recent *S. cerevisiae* transcriptome-wide ribosome profiling (Ribo-Seq) study failed to corroborate this result (Mangkalaphiban *et al*., 2021). Recent machine learning re-analysis of published Ribo-Seq datasets provided further support for the notion that the identity of P-site codon influences the readthrough efficiency, with AUA (isoleucine), CUG (leucine), and GAC (aspartate) as well as ACC and ACA (threonine) being associated with higher readthrough (Mangkalaphiban *et al*., 2024). In humans, the nature of the P-site codon was also shown to modulate the efficiency of readthrough, but the effect does not seem to be determined by the nature of -1 and -2 nucleotides, nor the amino acid encoded by the penultimate codon, but rather the nature of the P-site tRNA seems to be decisive (Loughran *et al*, 2023). As the ribosome-bound eRF1 interacts with the P-site tRNA, the nature of the P-site tRNA can also affect the stability of the termination complex (Skabkin *et al*, 2013), which could explain some P-site tRNA-specific effects.

In addition to the dedicated release factors eRF1 and eRF3, several other factors are known to influence translation termination. ABCE1/Rli1 not only promotes ribosome recycling, but also eRF1-mediated peptide release (Shoemaker & Green, 2011). Further, elongation factor eIF5A has critical roles in both elongation and termination (Pelechano & Alepuz, 2017; Schuller *et al*, 2017). Binding to the ribosomal E site, eIF5A accesses the peptidyl transferase center (PTC) to promote peptidyl transfer during elongation and eRF1-mediated peptide release during termination (Gutierrez *et al*, 2013; Pelechano & Alepuz, 2017; Rauscher *et al*, 2024; Schmidt *et al*, 2016; Schuller *et al*., 2017). Finally, the yeast elongation factor eEF3 prevents readthrough of premature stop codons by stimulating the esterase activity of the eRF1 (Kobayashi *et al*, 2023). A member of the ABCF ATPase protein family (Murina *et al*, 2019), eEF3 is an essential translation factor (Sandbaken *et al*, 1990) that promotes translocation step during protein synthesis by promoting the dissociation of deacylated tRNA from the ribosomal E site (Andersen *et al*, 2006; Ranjan *et al*, 2021).

We previously showed that *S. cerevisiae* ABCF ATPase New1 is involved in translation termination and/or ribosome recycling (Kasari *et al*, 2019b). Because the *new1Δ* yeast strain is cold-sensitive and displays a ribosome assembly defect, New1 was originally suggested to play a role in ribosome assembly (Li *et al*, 2009b). Using ribosome profiling, we demonstrated that the loss of New1 results in ribosomal queuing at stop codons and that this queuing is strongly associated with the presence of 3’-terminal lysine and arginine, and, to a lower extent, asparagine, codons (Kasari *et al*., 2019b). The effect was observed both at 20°C and 30°C, but it was stronger at 20°C. However, our Ribo-Seq experiments lacked the signal for the stop codon, and we were, therefore, unable to determine the effects on ribosome occupancy at the actual stop codons. The technical reason for the lack of the stop codon signal was likely the omission of cycloheximide during Ribo-Seq library preparation, as this antibiotic is known to stabilise post-termination complexes (Susorov *et al*, 2015), which are otherwise highly unstable (Schuller *et al*., 2017). Therefore, while accumulation of ribosomes at stop codons was implied based on the observed ribosomal queuing in the 3’ ORF region preceding the stop codon, the direct evidence for ribosomal accumulation at stop codons is lacking. Moreover, the relatively shallow coverage of our Ribo-Seq datasets meant that we were unable to establish if the inferred ribosomal stalling is a function of the C-terminal amino acid and/or if it required a specific mRNA codon. Finally, we also found that the overexpression of New1 can compensate for the loss of the essential translation elongation factor eEF3 (Kasari *et al*., 2019b), suggesting that New1 has a secondary function in elongation.

In this study, we have revisited the question of New1’s biological function using a combination of single-particle cryo-EM reconstructions, readthrough reporter assays, and 5PSeq. Our results confirm that, indeed, New1 is involved in the resolution of ribosomal stalling at stop codons preceded by an arginine or a lysine codon. Surprisingly, the stalling is exclusive to specific P-site tRNA isoacceptor species and is not defined solely by the nature of the C-terminal amino acid, as we have suggested earlier (Kasari *et al*., 2019b). Finally, we show that in addition to its role in translation termination, New1 samples the ribosome throughout its functional cycle, hinting at a more general role of New1 in translation.

## MATERIALS AND METHODS

### Yeast strains, plasmids, oligonucleotides, media, and genetic procedures

Yeast strains, plasmids and oligos used in this study are listed in **Supplementary Table 1**. Details of plasmid constructions are described in the relevant sections below. Yeast media was prepared as described (Amberg *et al*, 2005), with the difference that the composition of the drop-out mix was as per Johansson (Johansson, 2017). Difco Yeast Nitrogen base w/o amino acids was purchased from Becton Dickinson (291940), amino acids and other supplements from Sigma-Aldrich. YEPD medium supplemented with 200 µg/mL Geneticin (Gibco 11811-023) was used to select for cells harbouring the *kanMX6* marker (Longtine *et al*, 1998). YEPD medium supplemented with 100 μg/mL Nourseothricin clonNAT (Jena Bioscience AB102L) was used to select for cells containing the natMX6 marker.

Strain MJY1092 (*MATα ura3Δ0 leu2Δ0 his3Δ1 new1Δ::kanMX6*) was generated by transforming the diploid strain UMY2836 with a *new1::kanMX6* DNA fragment harbouring appropriate homologies to the *NEW1* locus. The *new1::kanMX6* DNA fragment was PCR amplified from pFA6a-*kanMX6* (Longtine *et al*., 1998) using primers oMJ489 and oMJ490. Following PCR confirmation, the generated heterozygous diploid was allowed to sporulate and MJY1092 was obtained from a tetrad. Strain VHY68, which is a *new1Δ::kanMX6* strain expressing the synthetic LexA-ER-haB112 transcription factor (Ottoz *et al*, 2014), was generated from a cross between MJY1092 and VHY61. A strain deleted for *the tR(CCU)J* gene was constructed by transforming a diploid strain, formed between MJY1170 (*MAT*a *ura3Δ0*) and MJY1171 (*MATα ura3Δ0*), with a *tr(ccu)jΔ*::*natMX6* DNA fragment. The DNA fragment was amplified from pAG25 (Goldstein & McCusker, 1999) using primers VHKT207 and VHKT208. The generated heterozygous strain was sporulated and strain VHY75 (*MAT*a *ura3Δ0 tr(ccu)jΔ::natMX6)* was obtained from a tetrad. Strains VHY87 (*MATα ura3Δ0 tr(ccu)jΔ::natMX6*) and VHY88 (*MATα ura3Δ0 tr(ccu)jΔ::natMX6 new1Δ::kanMX6*) were derived from a cross between VHY75 and MJY1173.

### Polysome profile analyses

Polysome profile analyses were performed essentially as described earlier (Kasari *et al*, 2019a).

### Preparation of 5PSeq libraries

Cultures of wild type (MJY1171), *new1Δ* (MJY1173), *tr(ccu)jΔ* (VHY87) *and new1Δ tr(ccu)jΔ* (VHY88) strains were grown in biological triplicates overnight at 20°C (or, alternatively, at 30°C), diluted to OD_600_ ≈ 0.05 in 50 mL of SC medium, and incubated in a shaking water bath (160 rpm) at 20°C (or 30°C) until OD_600_ ≈ 0.8. Cells were harvested by centrifugation for two minutes at 2000 rpm before freezing in liquid nitrogen. Total RNA was isolated from the frozen cell pellets by hot-phenol extraction as previously described (Johansson, 2017).

To prepare 5PSeq samples from New1-associated ribosomes, plasmid VHp911 harbouring New1 with a his_6_-TEV-FLAG3 (HTF) tag under the control of a β-estriadiol inducible promoter was constructed. Briefly, pRS316-insul-(lexA-box)4-PminCYC1 harbouring New1-TAP (VHp262) (Kasari *et al*., 2019b) was linearised using primers KTVH14 and KTVH15 and the HTF with homologous sequences to VHp262 was amplified with oligos KTVH16 and KTVH17 in order to replace the TAP tag with the HTF tag. Both fragments were then ligated using Gibson cloning to generate VHp911. 500 mL cultures of *S. cerevisiae* strain VHY68 carrying VHp911 were grown in SC-Ura medium in a baffled flask at 20°C with shaking at 160 rpm. Once the culture reached the OD_600_ of 0.3, the medium was supplemented with 2 µM β-estradiol to induce New1-HTF and grown for 4 hours. At OD_600_ ≈ 0.9, cells were harvested via vacuum filtration, frozen in liquid nitrogen and stored at –80°C until further use. New1 associated ribosomal complexes were then immunoprecipitated: cryomilled samples were resuspended in 10 ml of 1x Polymix buffer (95 mM KCl, 5 mM NH_4_Cl, 20 mM HEPES pH 7.5, 1 mM DTT, 5 mM Mg(OAc)_2_, 0.5 mM CaCl_2_, 8 mM putrescine, 1 mM spermidine supplemented with 0.5 mM ATP and 0.5 mM EDTA (Takada *et al*, 2020)). The lysate was cleared by centrifugation – 2000g, 10 minutes and incubated with 100 µl of equilibrated anti-FLAG M2 Affinity Gel for 2 hours with end over end mixing at 4°C for binding before loading to a gravity flow column. The column was then washed 10 times with 1x Polymix buffer before elution in 200 µl of 1x Polymix supplemented with 0.1 mg/mL FLAG3 peptide (Sigma). 200 µl of the input lysate was saved and incubated at 4°C to use as a control for 5PSeq analysis of pulldown samples. RNA from lysate and pull downs were extracted via extraction with hot phenol (Johansson, 2017). 5PSeq libraries were prepared as previously described (Zhang & Pelechano, 2021a).

### NGS and data analysis

Multiplexed 5Pseq libraries were sequenced for 51 cycles (single read) on an Illumina NextSeq 550 platform. The quality of Illumina reads was controlled using FastQC (Andrews, 2010). The adaptor sequence (5′-NNNNCTGTAGGCACCATCAAT-3′) was removed using Cutadapt (Martin, 2011). After removing reads mapping to non-coding RNA, reads were mapped to the *S. cerevisiae* reference genome R64-1-1.85 using STAR (Dobin *et al*, 2013) and UMI-tools (Smith *et al*, 2017) was used to exclude reads from PCR duplication. BedGraph profiles were produced from SAM files with SAMtools (Li *et al*, 2009a) and Plastid (Dunn & Weissman, 2016).

The analysis pipeline was implemented in Python3 and is available at GitHub repository (htttps://github.com/tmargus/5pSeq). Reads that mapped once to the genome were used for final analysis, the mapping of the 5’ position of reads was computed using fivePSeqMap.py. Mapped reads were normalised as reads per million (RPM). The ribosome queuing metric was computed using the Python compute_queuing_5PSeq_v3.py using default cut-offs. Gene coverage was calculated using HTSeq-count (Anders *et al*, 2015).

In the metagene profile analysis we took only genes meeting the following conditions: i) 3^rd^ quartile of CDS coverage on the selected window codons >0 (at least 75% of codons are covered), ii) total coverage of the CDS exceeded that of the UTRs. Then, for each gene and profiles’ position, counts were normalized relative to the coverage within a window spanning –100 to +100 nucleotides around the start and stop codons. Metagene plots were generated by calculating the median of these normalized values at each position. Polarity scores (quantitative metrics of ribosome distribution along the transcript length) were calculated using *papolarity* tool (Vorontsov *et al*, 2021). Coverage profiles were normalized using normalization factor and library size estimates from differential expression analysis separately for each bedGraph profile. Finally, we visualised coverage tracks using svist4get (Egorov *et al*, 2019).

### Dual-luciferase stop codon readthrough assays

In order to manipulate the sequence surrounding the stop codon of the dual-luciferase readthrough (DLR) plasmid pDB723 (Keeling *et al*, 2004), the *Bam*HI site upstream of the Renilla luciferase gene was mutated, resulting in plasmid VHp893 allowing easy insertion of sequences of interest into the *Sal*I and *Bam*HI cloning site in between the Renilla and Firefly luciferase reporter genes. To introduce the desired reporter sequences, complimentary oligos, including offset sequences (to generate sticky ends for ligation) were first phosphorylated using polynucleotide kinase (thermo) and then annealed by heating to 95°C for 5 mins and cooling to room temperature over 2 hours. Fragments were then ligated into VHp893 cut with Fast digest *Sal*I and *Bam*HI enzymes (Thermo). All reporter plasmid sequences and corresponding oligos are described in **Supplementary Table 1**.

The relevant strains carrying reporter plasmids based on VHp893, were grown overnight in SC-ura medium at 20°C or 30°C, diluted to OD_600_ :: 0.05 in the same medium, and grown until OD_600_ of ::0.5 at the respective temperature. 5 μL aliquots (in technical triplicates) from the cultures were assayed as previously described (Johansson & Jacobson, 2010). Readthrough assays were performed using the dual-luciferase reporter assay system (E1910, Promega) and a GloMax 20/20 luminometer (Promega). All experiments were performed at least three times using independent transformants for each experiment.

### Ex vivo *purification of New1p-80S ribosome complexes*

The New1p *ex vivo* pull-out was performed as described previously for eEF3 (Ranjan *et al*., 2021), but with a yeast strain (obtained from Horizon Discovery) expressing New1 C-terminally tagged with a tandem affinity purification (TAP) tag (derived from strain BY4741, genotype: *MAT*a *his3Δ1 leu2Δ0 met15Δ0 ura3Δ0*) (Ghaemmaghami *et al*, 2003). Briefly, cultures were grown to exponential phase and then harvested by centrifugation to pellet the cells. Cells were then lysed by glass bead disruption and incubated with IgG-coupled magnetic beads (Dynabeads M-270 Epoxy from Invitrogen) with slow tilt rotation for 1 h at 4°C in Buffer 30 (20 mM HEPES (pH 7.4), 100 mM KOAc, 10 mM Mg(OAc)_2_, 1 mM DTT). The beads were pelleted and washed three times using detergent containing Buffer 30 (supplemented with 0.05% TritonX), followed by an additional washing step in Buffer 30. The elution of the complex was done via cleavage of the TAP-tag by incubation with the AcTEV Protease (Invitrogen) for 3 h at 17 °C in Buffer 30.

### Sample and data collection

The eluate of the *ex vivo* New1p-80S complexes was crosslinked with 0.02% glutaraldehyde for 20 min on ice, and the reaction was subsequently quenched with 25 mM Tris-HCl (pH 7.5). The detergent n-Dodecyl-β-D-maltoside (Sigma, D5172) was added to the sample to a final concentration of 0.01% (v/v). For preparation of cryo-grids, 5 μL (8 A260/mL) of the freshly prepared crosslinked *ex vivo* New1p-80S complex was applied to precoated Quantifoil R3/3 holey carbon grids coated with 2 nm carbon and vitrified on a Vitrobot Mark IV (FEI, Netherlands), and stored in liquid nitrogen until use. Data collection was performed on a FEI Titan TEM microscope equipped with a Falcon2 detector. Data were collected at 300 keV with a total dose of 25 e^−^/Å^2^ fractionated over 10 frames with a pixel size of 1.084 Å/pixel and a target defocus range of −1.3 to −2.8 μm using the EPU software (Thermo Fisher).

### Cryo-EM data processing

Processing was performed using Relion 3.1 (Zivanov *et al*, 2018). The initial dataset consisted of 12,087 multi-frame movies. Movie frames were aligned with MotionCor2 using 5x5 patches (Zheng *et al*, 2017) and the CTF of the resulting micrographs was estimated using Gctf (Zhang, 2016). Only micrographs with an estimated resolution of 3.5 Å or better were retained (11,798 micrographs). From these, 381,667 particles were picked using crYOLO with the general model (Wagner *et al*, 2019; Wagner & Raunser, 2020). After 2D classification, 243,285 particles resembling ribosomes were selected and extracted with a box size of 140 pixels at a pixel size of 3.252 Å. These particles were aligned into an initial 3D volume using an empty 80S volume in the non-rotated state as a reference (PDB 4V88 (Ben-Shem *et al*, 2011)). From these aligned particles the following steps of 3D classification were performed without further angular sampling.

To select for New1-containing 80S ribosomes, a soft mask was designed around the known New1 binding site (Kasari *et al*., 2019b) and masked classification was performed into six classes. Only one class, consisting of 52,007 particles, contained density for New1 and was selected for further classification (= 100% New1). The next round of classification was performed without a mask into three classes which resulted in class1 with high resolution (32,711 particles, 63% of New1-containing particles, non-rotated), class2 with low resolution (11,889 particles, 23% of New1-containing particles, non-rotated) and class3 with high resolution (7,407 particles, 14% of New1-containing particles, rotated; designated STATE3). Class1 was selected for further classification without a mask and could be split up into two major states which showed density for either eIF5A or eRF1 (17,323 particles, 33% of New1-containing particles and 15,388 particles, 30% New1-containing particles, respectively).

For further analysis, partial signal subtraction was performed on these two classes using soft spherical masks around the peptidyl transferase center. The resulting particles were sorted using soft masks at the locations of eIF5A and eRF1, respectively. Each of the two could be further sorted into three classes. The eIF5A-containing particles separated into class1 (8,122 particles, 15% of New1-containing particles, A-tRNA/P-tRNA/eIF5a, designated STATE1), class2 (4,602 particles, 9% of New1-containing particles, A-tRNA/P/E*-tRNA, designated STATE2) and class3 (4,599 particles, 9% of New1-containing particles, A-tRNA/P-tRNA/E-tRNA, designated STATE5). The eRF1-containing particles separated into class1 (5,230 particles, 10% of New1-containing particles, P-tRNA/E-tRNA, designated STATE4), class2 (4,545 particles, 9% of New1-containing particles, A-tRNA/P-tRNA/E-tRNA, designated STATE6) and class3 (5,613 particles, 11% of New1-containing particles, eRF1/P-tRNA/E-tRNA, designated STATE7).

For high-resolution refinements, particles were re-extracted with a box size of 420 pixels at a pixel size of 1.084 Å. For CTF refinement only anisotropic magnification was refined. The pixel size of the final maps was estimated by comparison to existing structures using UCSF ChimeraX (Goddard *et al*, 2018; Pettersen *et al*, 2021). Local resolution estimation and subsequent filtering of volumes by local resolution were performed using Bsoft (Cardone *et al*, 2013; Heymann, 2018).

### Figure preparation

Molecular graphics were prepared with UCSF ChimeraX, developed by the Resource for Biocomputing, Visualization, and Informatics at the University of California, San Francisco, with support from National Institutes of Health R01-GM129325 and the Office of Cyber Infrastructure and Computational Biology, National Institute of Allergy and Infectious Diseases (Goddard *et al*., 2018; Pettersen *et al*., 2021). Figures were generated using ImageJ (Schneider *et al*, 2012), Adobe Photoshop, Adobe Illustrator and Inkscape.

### Quantification and statistical analysis of cryo-EM data

Relion uses a “molecular smoothness” prior within a regularized likelihood optimization framework when refining single-particle cryo-EM data (Scheres, 2012a, b). The “gold standard” FSC approach is used to prevent overfitting (Henderson *et al*, 2012; Scheres & Chen, 2012).

## RESULTS

### Cryo-EM structures of ex vivo New1-ribosome complexes

To ascertain the ribosome functional states that New1 can interact within the cell, we employed affinity chromatography in combination with *S. cerevisiae* cells expressing a chromosomally encoded TAP-tagged New1 protein (see *Materials and Methods*), as we performed previously for obtaining *ex vivo* eEF3-and Gcn-ribosome complexes (Pochopien *et al*, 2021; Ranjan *et al*., 2021). The New1-TAP eluate was stabilised through mild crosslinking with 0.02% glutaraldehyde treatment before being applied to cryo-EM grids and subjected to single-particle cryo-EM analysis. Focussed 3D classification revealed that 52,007 (21%) of the total 243,285 particles contained additional density within the New1 binding site (**Supplementary Figure 1**). Presumably either New1 dissociated from the other ribosomal particles during sample preparation and/or these ribosomes co-purified with New1-ribosome complexes because they were associated with the same mRNA in the polysomes. The putative New1-ribosome complexes were subjected to extensive 3D classification, yielding in total seven defined ribosome functional states, which we term states 1-7 (Figure 1A**-G** and **Supplementary Figure 1**). The distribution of particles between the states was similar, ranging from 9-15%, meaning that each substate was comprised of between 4,545-8,122 ribosome particles (**Supplementary Figure 1**). Despite the limited particle numbers, we were still able to obtain cryo-EM reconstructions of each state with average resolutions ranging from 5.1-6.0 Å (**Supplementary Figure 2A-G**). Although this resolution precludes a molecular interpretation, it was sufficient to assign the ligands and their positioning within the ribosomal complexes (Figure 1A-G). Six of the seven ribosome complexes (States 1-6) contained two or three tRNAs and were therefore assigned as elongating state ribosomes (Figure 1A-F), reminiscent of the many elongating complexes observed in the eEF3 pull-downs (Ranjan *et al*., 2021). In all these states, New1 spans the 40S head and central protuberance of the 60S subunit (Figure 1A-G), analogous to that observed previously for New1 in the *in vitro*-formed New1-80S complex (Kasari *et al*., 2019b) as well as for *in vitro* and *ex vivo* eEF3-ribosome complexes (Andersen *et al*., 2006; Pochopien *et al*., 2021; Ranjan *et al*., 2021). The seventh ribosome complex (State 7) contained A-and P-tRNAs as well as clear density within the A-site for eRF1 and was therefore considered a termination state (**Supplementary Figure 1**). State 7 appeared to be similar to the eRF1-eRF3-ribosome state observed in the Gcn20-pull downs (Pochopien *et al*., 2021).

**Figure 1.**
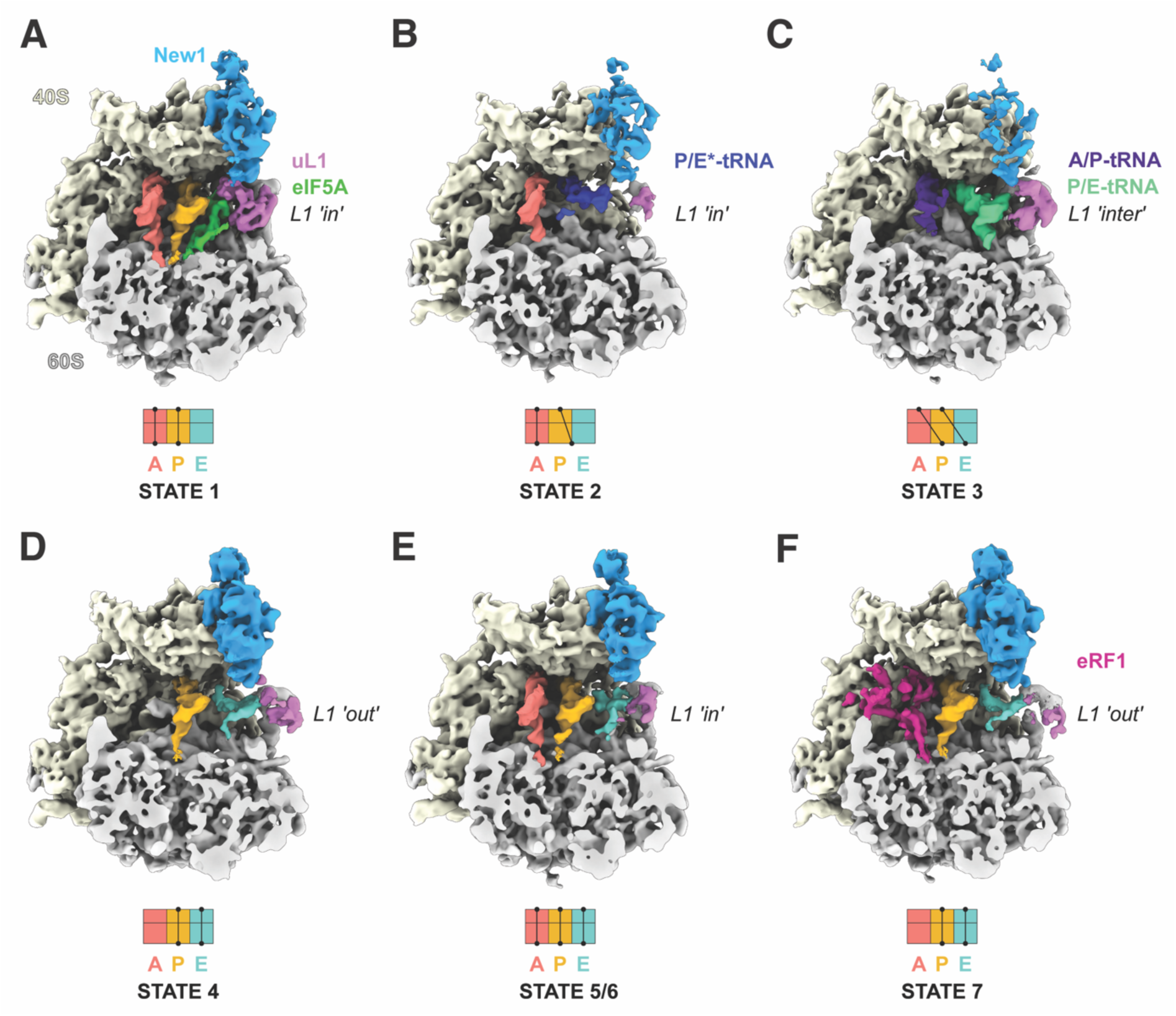
Cryo-EM structures of *ex vivo* New1-ribosome complexes. (A-F) cryo-EM maps of New1p (blue) in complex with (A-E) elongating (states 1-6) and (F) termination (State 7) state ribosomes. The 40S and 60S subunits are colored yellow and grey, respectively and the uL1 protein (pink) of the L1 stalk is indicated in the “in”, “out” or intermediate (“inter”) conformations. A-tRNA (red), P-tRNA (orange), E-tRNA (cyan), A/P-tRNA (dark blue), P/E-tRNA (green), P/E*-tRNA (dark blue), eIF-5A (lime) and eRF1 (magenta) are colored. The schematics below the maps indicate the conformation of the tRNAs in the (A-C) pre-translocational states (states 1-3) and (D-F) post-translocational states (States 4-7).

Of the elongating states, we observed New1 bound to three pre-translocation (PRE) and three post-translocation (POST) state ribosomes (Figure 1A-F). The canonical PRE state with A-and P-tRNAs (State 1) was also observed previously interacting with eEF3 (termed PRE-1; (Ranjan *et al*., 2021)), however, unlike in the case of eEF3, we observed additional density within the E-site that could be unambiguously assigned to the elongation factor eIF5A (Figure 1A). States 2 and 3 contained hybrid A/P-and P/E-tRNAs on non-rotated and rotated ribosomes, respectively (Figure 1B**, C**). New1 was poorly resolved in the rotated state, suggesting that it only binds stably to non-rotated ribosomes, as reported for eEF3 (Ranjan *et al*., 2021). State 4 represents a POST state with P-and E-site tRNAs, however, the L1 stalk adopts an outward conformation and the density for the E-site tRNA is poorly ordered (Figure 1D). We did not observe a POST state containing P-site tRNA but lacking E-site tRNA, as observed in the eEF3-pull downs (Ranjan *et al*., 2021). This would be consistent with the previous idea that New1 cannot efficiently complement eEF3 because of the shorter chromodomain that is less efficient at mediating E-site tRNA release (Kasari *et al*., 2019b). In this regard, it is interesting that tRNAs occupied all three tRNA binding sites (A-, P-and E-sites) in States 5 and 6 (Figure 1E**, F**), since this can only arise if E-site tRNA release is slower than A-site tRNA decoding and accommodation. States 5 and 6 appeared to differ from each other only with respect to the position of the L1 stalk (Figure 1E). Although three tRNAs on the ribosome were not observed in the eEF3 pull-downs, such functional states have been observed previously in *ex vivo*-derived human polysome samples (Behrmann *et al*, 2015). Collectively, the ensemble of New1-ribosome structures suggests that New1 can interact with both elongating and terminating ribosomes and may preferentially favour ribosomes that encounter obstacles leading to recruitment of eIF5A or accumulation of three tRNAs per ribosome.

### New1 samples the ribosome throughout the translation cycle

Given that our Ribo-Seq analysis specifically implicated New1 in translation termination and/or ribosome recycling (Kasari *et al*., 2019b), the cryo-EM results showing its association with elongating ribosomes are surprising. Therefore, we next used 5PSeq to map the positions of immunopurified New1-associated ribosomes transcriptome-wise. 5PSeq is functional genomics method that is highly complementary to Ribo-Seq (Pelechano *et al*, 2015). To infer ribosomal positions on mRNA, 5PSeq relies on specifically sequencing 5’-phosphorylated mRNA fragments that are generated during co-translational 5’-3’ mRNA decay, and the resulting mRNA 5’ ends represent the borders of mRNA protected by the ribosome, thus allowing mapping of ribosomes under a given condition (Pelechano *et al*., 2015; Pelechano *et al*, 2016). Since the method relies on the 5’-3’ exonucleases “catching up” with stalled ribosomes, it is particularly well-suited for studying defects in intrinsically slow translational events, such as termination (Pelechano & Alepuz, 2017; Pelechano *et al*., 2016; Zhang & Pelechano, 2021b). A HTF-tagged New1 was ectopically expressed for 2 hours in yeast cells grown at 20°C followed by cell disruption and immunopurification. The total mRNA for 5PSeq library preparation was extracted from both the pulldown samples as well as from the input lysate (**Supplemental Figure 3**). The latter sample served as a control representing the global cellular translation. Note that, just as in the case of *ex vivo* New1-ribosome cryo-EM reconstructions, a possible caveat is that a fraction of the 5PSeq signal in the New1 pulldown samples could be coming from ribosomes that are not directly associated with the factor but, rather, co-purify via a shared mRNA. However, regardless this potential limitation, the pulldown sample is strongly enriched in New1-associated ribosomes, thus allowing a meaningful comparison with the lysate-derived 5PSeq dataset.

The metagene analysis of our 5PSeq experiments revealed a striking similarity between the lysate (black) and New1 pulldown (blue) libraries (Figure 2A-B). Both samples display excellent periodicity within the ORF body, as well as near-identical peaks at -14 nt relative to start codons (corresponding to a start codon positioned in the ribosomal P-site) (Figure 2A) and at -17 nt relative to stop codons (corresponding to stop codons positioned in the A-site) (Figure 2B) (Pelechano *et al*., 2015, 2016). We calculated the polarity score, a metric that quantifies the ribosomal distribution along the ORF (Schuller *et al*., 2017), for the two datasets. Distributions of polarity scores for New1-puldown and lysate-derived samples were near-identical (Figure 2C), indicative of New1 sampling the ribosomes throughout the translation cycle. Finally, we asked the question whether New1 is associated with terminating ribosomes preferentially stalled with arginine or lysine codons in the P-site. Note that as the experiments were performed with a yeast strain expressing functional New1, we did not expect to observe a dramatic defect that we observed in the *new1Δ* strain (Kasari *et al*., 2019b). To quantify the extent of ribosomal stalling for individual ORFs, we applied the queuing score (QS) metric that quantifies the extent of the ribosomal “pile-up” in front of stop codons (Kasari *et al*., 2019b). Both in the New1 pulldown and lysate-derived libraries, arginine-encoding C-terminal codons displayed the highest QS, suggestive of relatively inefficient termination (Figure 2D). However, the extent of stalling was similar for the lysate and in New1-pulldown samples, signifying a lack of specific enrichment. Taken together, in good agreement with the cryo-EM results, our 5PSeq data suggest that New1 samples ribosomes throughout the translational cycle.

**Figure 2.**
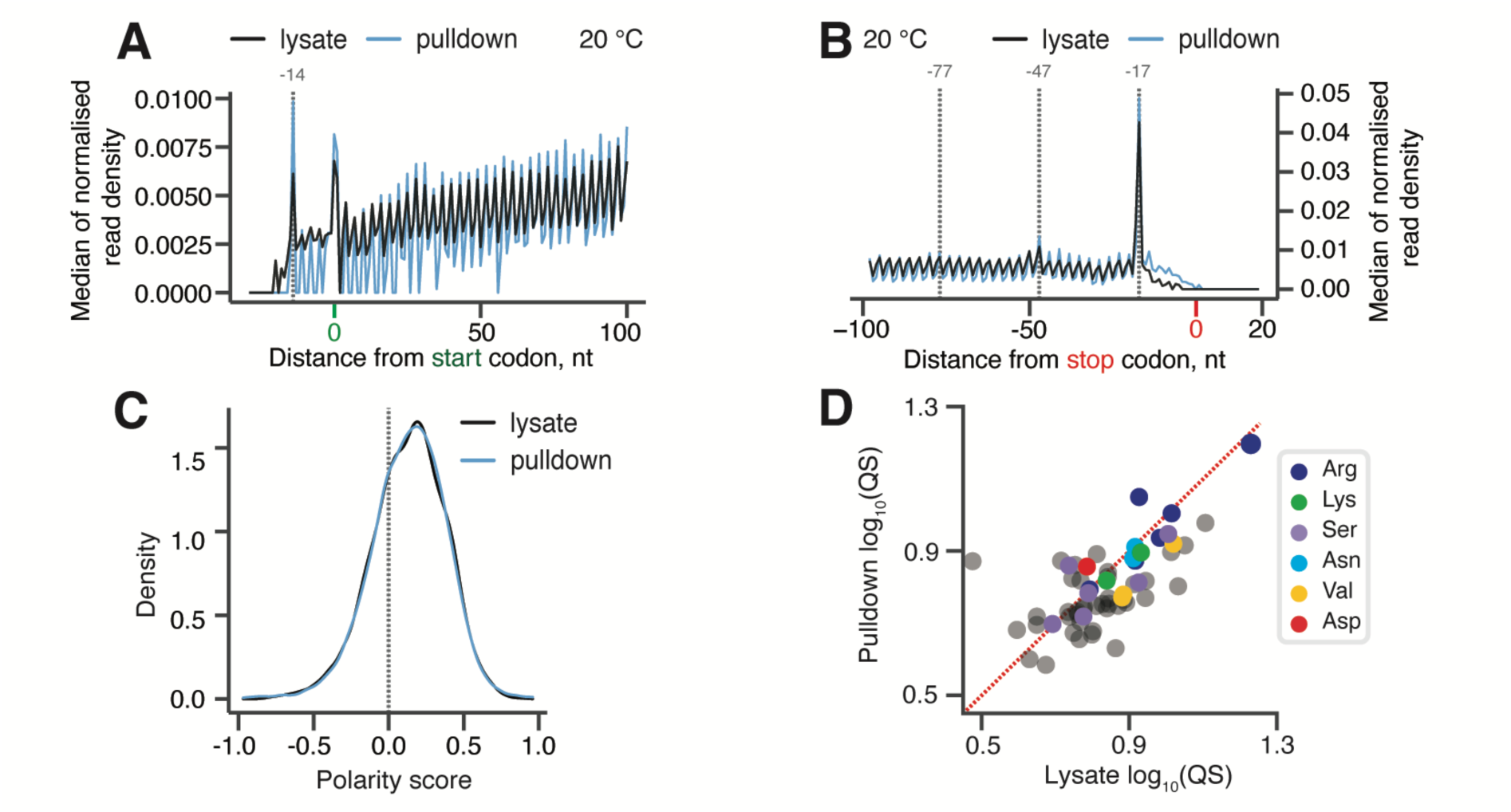
New1 samples the ribosome throughout the translation cycle. (**A, B**) 5PSeq metagene analysis. The 5’ read ends for of pulldown (blue) and lysate (black) samples aligned to start (A) and stop codons (B). Stalls represent the 5’ boundary of the ribosome-protected 5’-phosphorylated mRNA fragment. The 5’ position is offset by -14 nt from the fist nucleotide in the P site and -17 nt from the first nucleotide in the ribosomal A site. (**C**) Distributions of ribosome polarity scores across transcriptome for immunoprecipitated New1-HTF ribosome complexes (pulldown, blue) and the input lysate (black). Positive and negative scores reflect enrichment of reads towards either 3’ and 5’ end of CDS, respectively. (**D**) Ribosomal queuing scores (QS) calculated for ORFs parsed by the codon preceding the stop codon. The red dashed line is a guide for the expected position of data points in the absence of the systematic difference between in QS between the pulldown and lysate samples. All analyses were performed on pooled lysate and New1 pulldown datasets from four biological replicates each.

### 5PSeq reveals that the lack of New1 is associated with ribosomal stalling at stop codons preceded by an arginine, lysine or asparagine codon

To uncover the effect of New1 loss on ribosomal stalling genome-wide, we performed a 5PSeq analysis of wild-type and isogenic *new1Δ* yeast strains, both grown at 20°C as the ribosomal “pile-up” in front of the stop codon observed by Ribo-Seq was stronger at this temperature (Kasari *et al*., 2019b). The 5PSeq coverage for individual ORFs was strongly correlated for individual replicates (R^2^ = 0.957 and 0.961 for wild-type and *new1Δ*, respectively, **Supplementary Figure 4A,B**), thus allowing us to use pooled data for the analysis. Metagene analysis of the wild-type dataset readily detected a pronounced peak at the position -17 from the stop codon, which corresponds to terminating ribosomes (Pelechano *et al*., 2015) (see Figure 3A, black trace, for pooled 5Pseq data for all three individual replicates, and **Supplementary Figure 4C** for individual replicates analysed separately). In the *new1Δ* 5PSeq metagene plots, the corresponding peak is 1.7-fold higher, and is preceded by additional peaks spaced at 30 nucleotide intervals (Figure 3A, red trace, and **Supplementary Figure 4D**); note that these additional peaks were also detected by Ribo-Seq analysis (Kasari *et al*., 2019b).

**Figure 3.**
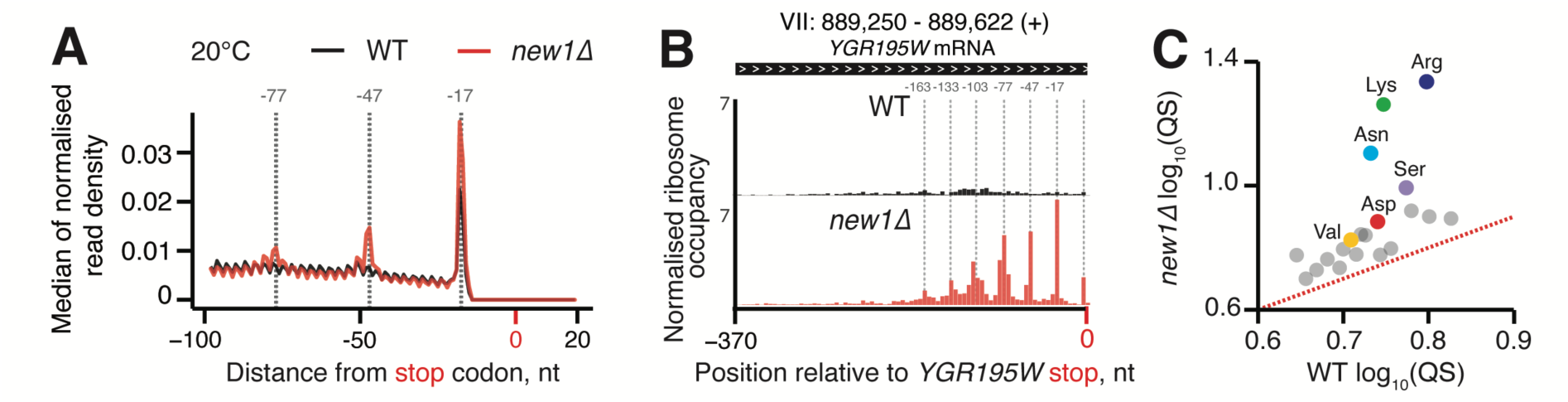
New1 loss is associated with ribosomal queuing upstream of stop codons proceeded by C-terminal arginine, lysine and asparagine. (**A**) Metagene analysis of the 5’ read ends for the 5PSeq libraries generated from wild-type (WT MYJ1171, black trace) and New1-deficient (*new1Δ* MYJ1173, red trace) yeast strains grown at 20°C. Stalls represent the 5’ boundary of the ribosome-protected 5’-phosphorylated mRNA fragment. The 5’ position is offset by -17 nt from the first nucleotide in the ribosomal A site. With an 80S ribosome covering 30 mRNA bases, the 30-nt periodicity of the in the *new1Δ* metagene is indicative of multiple ribosomes queued at the stop codon. (**B**) An example of a heavily “queued” ORF *YGR195W* encoding a C-terminal AGG arginine codon. Normalised read coverage around the stop codon is shown. (**C**) Ribosomal queuing scores (QS) calculated for ORFs parsed by the codon preceding the stop codon. The red dashed line is a guide for the expected position of data points in the absence of the systematic difference between in QS between the wild-type and *new1Δ* samples. All analyses were performed on pooled wild-type and *new1Δ* 5PSeq datasets from three biological replicates each.

To quantify the extent of ribosomal stalling at the stop codon for individual ORFs we next applied the QS metric (**Supplementary Table 2**). In the case of individual ORFs with high QS, up to seven peaks corresponding to ribosomes stalled at the stop codon were clearly detectable, as exemplified by a highly queued ORF of *YGR195W* that ends with C-terminal AGG arginine codon and encodes SKI6, a 3’-5’ RNase (Figure 3B). In the case of the wild-type strain, we detect no correlation between the QS of individual ORFs in 5PSeq and Ribo-Seq datasets (**Supplementary Figure 4E**), which is likely due to the inability to detect the stop codon peak in the Ribo-Seq dataset as well as with the lack of queued ribosomes in the wild-type strain. Conversely, queuing scores of individual ORFs of the *new1Δ* 5PSeq dataset are positively correlated (R = 0.4668) with that for the Ribo-Seq dataset (**Supplementary Figure 4F**). This demonstrates that despite 5PSeq and Ribo-Seq relying on two different approaches to detect ribosomal stalling (posttranslational mRNA degradation and mRNA protection from RNA digestion by the ribosomes, respectively), the two experiments detect an overlapping set of ORF-specific signals. Importantly, the fold change in QS between wild-type and *new1*Δ strains for each ORF is also correlated in both datasets (R = 0.4336, **Supplementary Figure 4G**). However, QS tended to be higher in the Ribo-Seq dataset, which is to be expected, since 5Pseq produces a read corresponding to only the most 5’ ribosome on the mRNA, and thus does not detect multiple ribosomes in multi-ribosome “pile-ups’. Finally, we have computed QS for the 5PSeq dataset sorted by the nature of C-terminal amino acid. In excellent agreement with the Ribo-Seq analysis (Kasari *et al*., 2019b), C-terminal arginine (R), lysine (K) and, to a lower extent, asparagine (N) are associated with a high degree of ribosomal queuing at stop codons in the *new1Δ* strain (Figure 3C).

### The stop codon identity alone does not determine the extent of ribosomal queuing in new1Δ cells

Since the nature of the stop codon is the key determinant of termination efficiency (Bonetti *et al*., 1995), we next plotted the distributions of the queuing scores for individual ORFs with 5PSeq data parsed by the stop codon identity. Distributions for wild-type and *new1Δ* datasets are similar, with the *new1Δ* distribution slightly shifted towards a higher queuing score (Figure 4A-C). The median queuing scores for all the three stop codons were similar both in wild-type (UAA 0.71, UAG 0.72 and UGA 0.76) and *new1Δ* (UAA 0.90, UAG 0.93 and UGA 0.94) datasets, displaying 1.52-1.62-fold increase upon New1 loss. Next, focusing on the stop codon with lowest fidelity, UGA, we analysed the effect of the 3’ +4 nucleotide of the extended stop codon. Of the four possible extended stop codons, UGA followed by a C has the lowest fidelity (Bonetti *et al*., 1995). While we observe slightly higher queuing score in the case of this extended stop codon, the effect is similar in both wild-type (UGA-G 0.77, UGA-A 0.77, UGA-U 0.73 and UGA-C 0.83) and *new1Δ* (UGA-G 0.91, UGA-A 0.93, UGA-U 0.94 and UGA-C 1.10) strains (**Supplementary Figure 5A-F**). Collectively, these results demonstrate that the nature of the stop codon alone has a relatively weak effect on the extent of the ribosomal queuing in the *new1Δ* strain at the stop codon.

**Figure 4.**
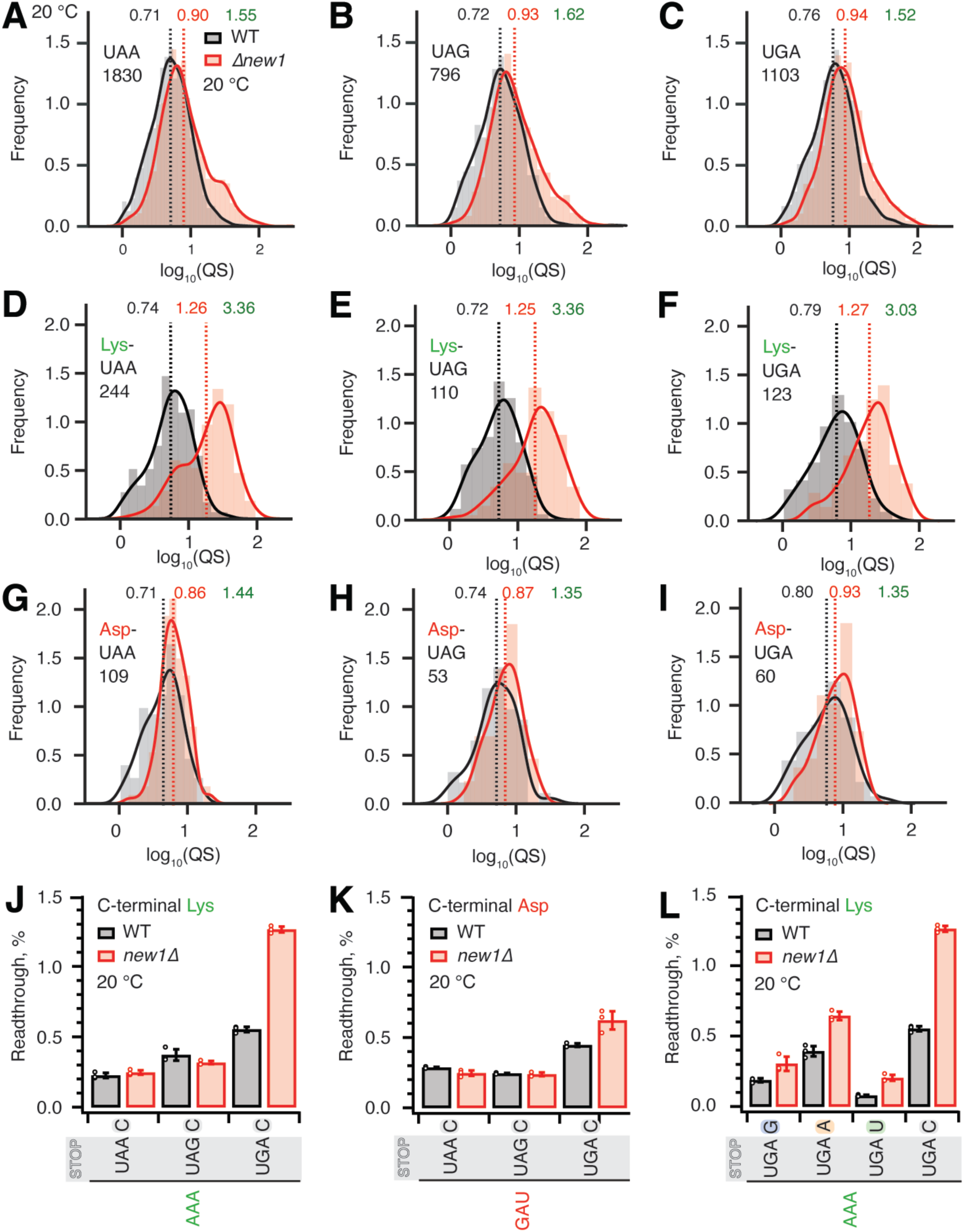
The absence of New1 specifically increases ribosomal readthrough when the weak stop codon UGA is preceded by C-terminal lysine. (**A**-**C**) Binned distributions of ribosome queuing scores (QS) for ORFs terminating in either UAA (A), UAG (B) or UGA (C) stop codons. (**D-F**) Same as (A-C), but with for ORFs encoding C-terminal lysine residues. Geomeans of QS distributions for wild type is black and for *new1Δ* is in red, the QS fold change is in green. Numbers of ORFs included in the analyses are shown on the figure, e.g. 1830 instances of ORFs with UAA stop codon. All analyses were performed on pooled datasets from three replicates collected at 20°C. (**G-H**) Same as (D-F), but with for ORFs encoding C-terminal aspartic acid. (**J**,**K**) Stop codon readthrough efficiencies in wild-type and *new1Δ* strains measured with dual-luciferase reporters harbouring UAA, UAG and UGA stop codons in combination with either a C-terminal AAA lysine codon (J) or GAU aspartic acid codon (K). (**L**) Readthrough efficiency of UAG G, UAG A, UAG U and UAG C extended stop codons in combination with a C-terminal AAA lysine codon in both wild-type and *new1*11 strains. Note that the same AAA UGA C results are shown on both panel (J) and (L) in order to make the data comparison easier. Error bars represent standard deviations. All experiments were performed at 20°C.

### New1 loss results in increased readthrough of weak extended stop codon UGA C preceded by the common AAA lysine codon

Next, we further parsed our datasets by the nature of the C-terminal amino acid and extracted the data for ORFs ending with lysine (associated with a strong ribosomal queuing in *new1Δ*) or asparagine (associated with a weak ribosomal queuing in *new1Δ*). For all the three stop codons preceded by lysine, we observed dramatically increased queuing in *new1Δ* (Figure 4D-F, **Supplementary Figure 6A-C**), while in the case of asparagine, as expected, the queuing scores are similar between the wild-type and *new1Δ* datasets (Figure 4G-H, **Supplementary Figure 6D-F**). In both cases the QS distributions are largely insensitive to the nature of the stop codon.

While we observed similar extents of ribosomal queuing in *new1Δ* at the three stop codons, the effect of New1 loss on efficiency of translation termination could still be sensitive to the nature of the stop codon, i.e. the same degree of queuing could have different mechanistic implications depending on the stop codon fidelity. Using dual-luciferase readthrough assays to probe the translation termination efficiency in yeast strains grown at 20°C, we tested the effects of C-terminal lysine (AAA) and C-terminal aspartic acid (GAU, the non-queued control) on readthrough of UAA, UAG and UGA codons followed by a 3’ +4 C (Figure 4J**,K**). In good agreement with previous reports (Bonetti *et al*., 1995; Cridge *et al*., 2018; Urakov *et al*, 2017), UGA is associated with the highest degree of readthrough in both wild-type and *new1Δ* cells. In the presence of C-terminal lysine the UGA readthrough is 2.3-fold higher in *new1Δ* (Figure 4J), while we detected a mere 1.4-fold increase in the case of the non-queued C-terminal aspartate (Figure 4K). We have also tested the effects of the +4 nucleotide following AAA UGA (Figure 4L). The loss of New1 results in increased readthrough in all extended stop reporters, with the effect ranging from 1.6 to 2.6-fold. In good agreement with (Bonetti *et al*., 1995), UGA C has the highest readthrough regardless of strain.

### Increased ribosomal queuing and stop codon readthrough in new1Δ cells are associated with a sub-set of penultimate codons

The high coverage of our 5PSeq dataset and strong 5PSeq signal at stop codons allowed further parsing of the dataset for individual codons encoding the C-terminal amino acid. Since the queuing score is only very moderately affected by the nature of the stop codon (Figure 4A-C), we next calculated the queuing scores for individual ORFs for data parsed by individual C-terminal codons, irrespective of the stop codon. Strikingly, individual P-site codons of the termination complex that encode the same amino acid are associated with dramatically different queuing scores (Figure 5A and **Supplementary Table 3**). While the penultimate AGG arginine codon is associated with increased ribosomal queuing in *new1Δ* cells, AGA is not. Similarly, while the penultimate AAA lysine codon is queuing-prone, AAG is not. Importantly, both AAA and AAG are common codons, suggesting that codon rarity, as such, is not a decisive factor. We plotted queuing score distributions for individual ORFs grouped by the nature of the individual codons encoding C-terminal amino acids: lysine (AAG and AAA, Figure 5B**,C**) and arginine (AGA and AGG, Figure 5D**,E**). The results are in agreement with the scatter plot of median queuing scores (Figure 5A).

**Figure 5.**
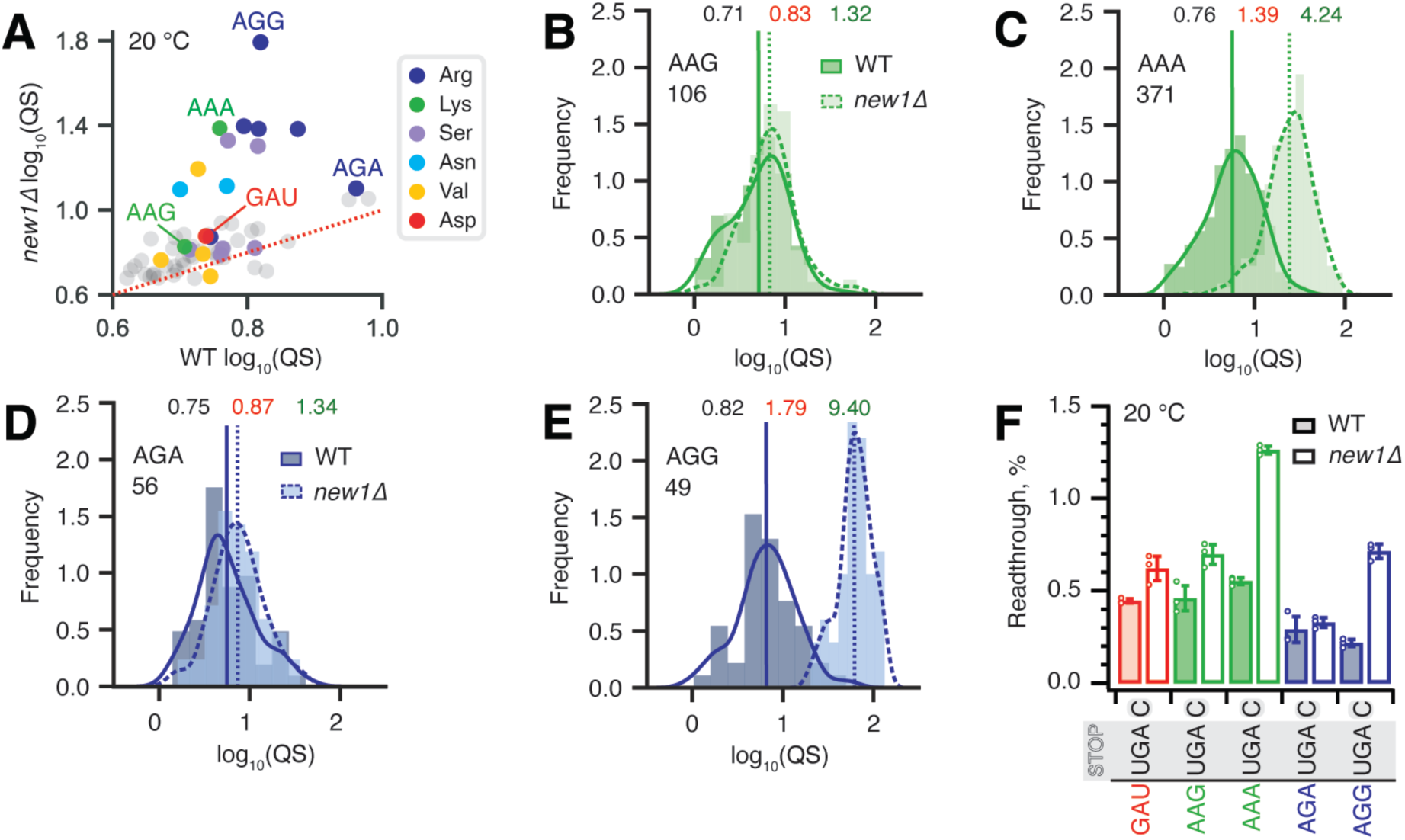
Synonymous codons encoding the C-terminal lysine and arginine residues have dramatically different effects on translation termination defects in *new1Δ* yeast. (**A**) Wild-type and *new1Δ* QS scores for ORFs parsed by the codon preceding the stop codon. (**B-E**) QS distributions with ORFs containing C-terminal lysine encoded by either AAG (B) or AAA (C), as well as arginine encoded by either AGA (D) or AGG (E). Wild-type: solid line, darker shading. Δ*new1*: dashed line, lighter shading. Geomeans of QS distributions for wild-type is black and for *new1Δ* is in red, the QS fold change is in green. All analyses were performed on pooled dataset from three replicates collected at 20°C. (**F**) Stop codon readthrough efficiencies of UGA stop codon combined with penultimate codons as indicated on the figure. Error bars represent standard deviations, all experiments were performed at 20°C.

We then tested the functional significance of the codon-specific ribosomal queuing in *new1Δ* using readthrough assays (Figure 5F). In agreement with the 5PSeq data, New1 loss results in increased readthrough in the case of AAA (but not AAG) as well as AGG (but not AGA). No apparent effect on readthrough was observed for the non-queued control GAU codon encoding aspartic acid (Figure 5F) (**Supplementary Figure 6**). Collectively, our results establish that the nature of the codon preceding the stop codon, and not just the C-terminal amino acid encoded by the said codon, is the crucial determinant for the translation termination defect observed in the absence of New1.

### Specific P-site tRNA isoacceptors are associated with increased readthrough of UGA stop codons in the new1Δ strain

In principle, the P-site codon-specific termination defect in *new1Δ* cells could be due to the nature of the mRNA codon located in the P-site, the P-site tRNA itself or a combination of both. A possible explanation could be that the rarity of the codon plays the decisive role, however, the queuing-prone nature of the common AAA lysine codon speaks against this hypothesis. Alternatively, specific tRNA isoacceptors reading the P-site codon could be responsible for the effect. To investigate this hypothesis, we parsed the ribosomal queuing scores calculated for 5PSeq data parsed on the basis of the individual P-site codons preceding the stop signal (**Supplementary Table 3**), correlated the parsed data with the *S. cerevisiae* isoacceptor tRNA species that decode the respective codons (Johansson *et al*, 2008), and identified the codons with increased (> two-fold) ribosomal stalling (Figure 6A, green boxes). These codons include the valine codon GUG (but not GUU, GUC or GUA), the Asn codons AAU and AAC, the lysine codon AAA (but not AAG), the serine codons AGU and AGC (but not the UCN codons) as well as the arginine codon AGA (but not AGG). Genetic manipulations of the tRNA species decoding these codons are complicated by the fact that they are often encoded by multiple genes, e.g. the cognate tRNA isoacceptor for the Lys AAA and AAG codons are encoded by 7 and 14 genes, respectively. However, the distribution of genes for the tRNA species decoding the arginine AGG (queued) and AGA (non-queued) codons provides an experimentally amenable system. The AGG codon is decoded by two tRNA isoacceptor species: the cognate rare 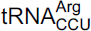 (encoded by a single gene, *tR(CCU)J*; decodes exclusively AGG codons) and the near-cognate, more abundant 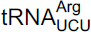 (which is encoded by 11 genes; able to decode both the AGG and AGA codon) (Johansson *et al*., 2008). In *new1Δ* cells, a P-site AGG codon is associated with ribosomal stalling at the stop codon, whereas an AGA codon is not. A possible mechanistic explanation is that the presence of P-site 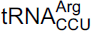 in pre-termination complexes results in inefficient termination which is, in turn, compensated by the action of New1.

**Figure 6.**
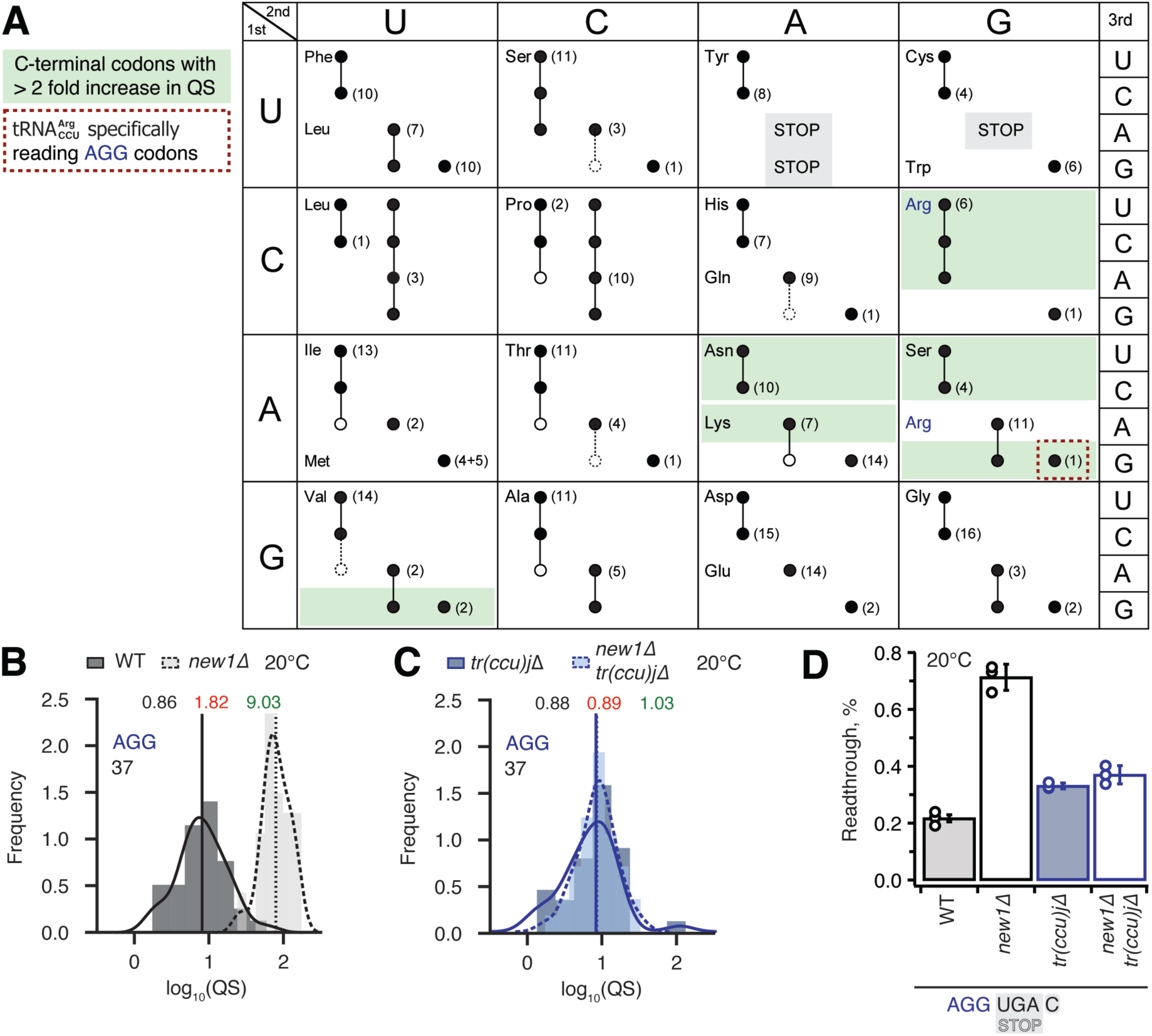
Termination defect in *new1Δ* yeast is dependent on the nature of the tRNA isoacceptor species present in the P-site of the pre-termination complex. (A) Decoding abilities of tRNA isoacceptors, adapted from Johansson and colleagues (Johansson *et al*., 2008). Codons read by individual tRNA species are indicated by circles connected by lines. The gene copy number for each tRNA species is indicated and the position of the number denotes the cognate codon. Empty circles indicate that the tRNA is less likely to read the codon. The empty dashed circles indicate that the tRNA does not efficiently read the codon. C-terminal codons that display increased QS in the *new1Δ* strain are highlighted with green shading. (B,C) Queuing score distributions for *NEW1* (solid outline) and *new1Δ* (dashed) strains either expressingor lacking (C) 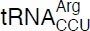. (D) Penultimate AGG codons are not associated with readthrough in *new1Δ* yeast when decoded by the near-cognate 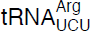 species. Readthrough efficiency for UGA stop codon preceded by penultimate AGG codon in strains either expressing or lacking 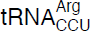. All experiments were performed at 20 °C.

Therefore, we decided to strictly allow only the near-cognate 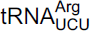 to decode AGG codons in the absence of competition with the 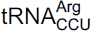 cognate 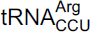. If the termination defect in *new1Δ* cells is induced by the P-site 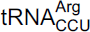, the phenotype should be suppressed by deleting the non-essential and single copy *tR(CCU)J* gene. To test this prediction, we constructed wild-type and *new1Δ* strains deleted for the *tR(CCU)J* gene and performed 5PSeq. The lack of [inlilne] in *new1Δ* cells reduced the ribosomal queuing at stop codons proceeded by AGG to near wild-type levels (*new1Δ* = 1.83 and *new1Δ tr(ccu)jΔ* = 0.89; note that the QS is calculated as log_10_). At the same time, the levels of ribosomal queuing at these stop codons are similar in the wild-type (0.86) and *tr(ccu)jΔ* (0.89) strains (Figure 6B**,C**). Finally, we performed readthrough assays in the wild-type, *new1Δ, tr(ccu)jΔ* and *new1Δ tr(ccu)jΔ* strains. Directly supporting our hypothesis, the increased readthrough in *new1Δ* cells was lost upon deletion of *tR(CCU)J*, suggesting that the presence of AGG-decoding 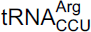 in the P-site could be, indeed, responsible for the termination defect (Figure 6D). Collectively, these results support a model where New1 is an accessory termination factor that assists termination on weak stop codons when specific isoacceptor tRNAs are present in the P-site.

### Ribosomal queuing and termination defects of new1Δ strain are uncoupled from the growth defect at low temperature

While the *new1Δ* strain grown at 20°C displays both growth and ribosome assembly defects (Li *et al*., 2009b), at 30°C the growth defect is negligible (Kasari *et al*., 2019b). Our analyses of polysome profiles confirm a reduction in the 40S:60S ratio in the *new1Δ* strain at 20°C, and, whilst this strength of the effect is reduced, the defect is still present at 30°C (Figure 7A,**B**). Importantly, our Ribo-Seq experiments were performed both at 20°C and 30°C, and, while the effect was stronger at 20°C, the ribosomal “pile-up” in front of the stop codon was also observed at 30°C (Kasari *et al*., 2019b).

**Figure 7.**
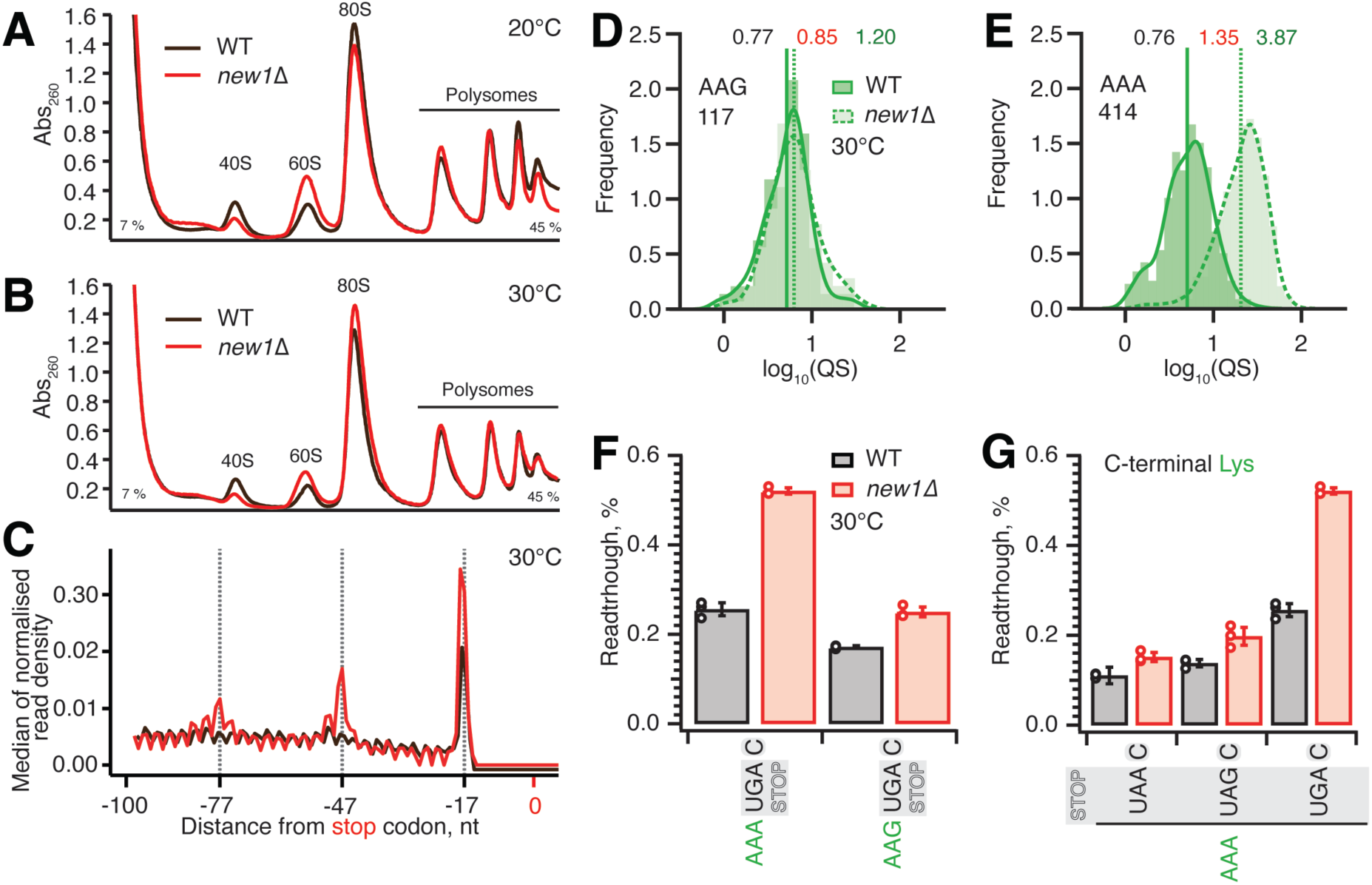
Ribosomal queuing and termination defects of *new1Δ* strain are uncoupled from the growth defect at low temperature. (**A, B**) Polysome profile analysis of wild-type (MYJ1171) and *new1Δ* (MYJ1173) yeast cells grown exponentially at 20°C (F) or 30°C (G) in SC media. Polysomes were resolved on 7-45 % sucrose gradients. All experiments performed at 30°C with the exception of (F), 5PSeq data analysis was performed on pooled data sets from three biological replicates. (**C**) Metagene analysis of the 5’ read ends for the 5PSeq libraries generated from wild-type (WT MYJ1171, black trace) and New1-deficient (*new1Δ* MYJ1173, red trace) yeast strains grown at 30°C. (**D**,**E**) QS distributions sorted by AAG lysine (D) or AAA lysine C-terminal codons (E), geomean QS values in the wild-type and *new1Δ* strains are given in black and red respectively and fold change in QS in green. (**F**) Readthrough of C-terminal AAA and AAG lysine codons in combination with a UGA stop codon. Error bars represent standard deviations. (**G**) Readthrough values of UAA, UAG and UGA stop codons in combination with a C-terminal AAA lysine codon. Error bars represent standard deviations. To make the data comparison easier, the same AAA UGA C results are shown in panel (F) and (G).

To test the causative connection between the translation termination effect caused by the New1 and cold-sensitivity, we have performed 5PSeq analysis as well as polysome profile analysis of wild-type and *new1Δ* strains grown at 30°C. In agreement with Ribo-Seq performed at 30°C (Kasari *et al*., 2019b), the metagene plots of 5PSeq datasets reveal New1-dependant queuing at 30°C is present, although reduced, when compared to the 20°C datasets (Figure 7C). Furthermore, like the 20°C datasets, this queuing is dependent on the nature of the C-terminal codon, as illustrated by AAG (non-queued) and AAA (queued) lysine codons which show a 1.2-fold and 3.9-fold increase in the *new1Δ* strain respectively (Figure 7D,**E**). We have also performed a corresponding set of readthrough assays at 30°C. While the readthrough efficiency is between two-and three-fold lower at 30°C than at 20°C, the effect of the *new1Δ* allele is essentially the same at 30°C: *new1Δ* cells show increased readthrough when the UGA but not the UAA or UAG stop codon is preceded by the AAA lysine codon (Figure 7F,**G**). Collectively, our results demonstrate that, irrespective of the strength of the temperature-dependent growth defect, increased ribosomal queuing at the stop codon in New1-deficient yeast correlates with increased readthrough of the low-fidelity UGA codon.

## DISCUSSION

Our 5PSeq and cryo-EM analyses of *ex vivo* New1-80S pulldown samples demonstrated that New1 interacts with both elongating and terminating ribosomes. As we have shown earlier, overexpression of New1 can compensate for the loss of otherwise essential elongation factor eEF3 (Kasari *et al*., 2019b), indicative of the functional overlap between the two ABCFs ATPases. However, eEF3 overexpression does not compensate for the loss of New1, consistent with a New1-specific function in translation termination and/or ribosome recycling (Kasari *et al*., 2019b).

By comparing the wild-type and *new1Δ* 5Pseq datasets, we identified the sequence determinants associated with translation defects on specific mRNA that occur in the absence of New1. Using dual-luciferase readthrough assays, we then validated the stop codon-and context-specific defects of translation termination in *new1Δ* cells. We demonstrate that in the absence of New1, translation termination efficiency is affected by the nature of the P-site codon / P-site tRNA isoacceptor tRNA decoding C-terminal arginine and lysine amino acids. While a AAA lysine codon 5’ of the stop codon is strongly associated with ribosome stalling and increased readthrough of the weak UGA stop codon, an AAG lysine codon is not. A similar behaviour was observed for the AGG / AGA arginine codon pair. Our results explain why in our previous study we failed to detect any effect of New1 loss when *YDR099W* / *BMH2* and *YNL247* 3’ ORF regions were tested in readthrough assays (Kasari *et al*., 2019b): while the AAA lysine codon preceding the stop codon in these reporters is, as we show here, associated with strong ribosomal queuing, due to high-fidelity UAA and UAG stop codons one would not expect increased readthrough in *new1Δ* cells. Collectively, our results suggest that by fine-tuning the termination efficiency, New1 suppresses and efficiently masks codon/tRNA-specific variation in termination efficiency in *S. cerevisiae*.

A comparison of the sequences and modifications of the different tRNA isoacceptors that cause ribosomal stalling in the absence of New1 did not reveal any obvious pattern to explain the phenomenon. However, structures of eukaryotic termination complexes reveal that the peptidyl-tRNA bound in the P-site directly interacts with eRF1 bound in the A site (Brown *et al*., 2015; Coelho *et al*, 2024). Specifically, the N-domain of eRF1 (involved in stop codon recognition) can form direct hydrogen bond interactions with the backbone of nucleotides located in the anticodon stem-loop of the P-site tRNA (**Supplementary Fig. 7A, B**). Thus, one can envisage that differences in the sequence and modifications of the tRNA isoacceptor located in the P site could directly influence the efficiency of the eRF1 functions during termination. Unfortunately, the only available structures of eukaryotic termination complexes are mammalian, nevertheless they are very consistent with our cryo-EM density for State 7 of the New1-bound termination complex (**Supplementary Fig. 7C**). We note that these complexes do not have the relevant tRNAs in the P site; for example, the highest resolution termination complex is from rabbit and has tRNA^Val^ in the P site rather than yeast tRNA^Arg^ or tRNA^Lys^ (Coelho *et al*., 2024). Moreover, while there are structures of ribosomes stalled with AAA-decoding 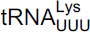 in the P site (Chandrasekaran *et al*, 2019; Tesina *et al*, 2020), there are no structures with 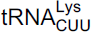 to which to compare them. *In silico* modelling of AAA-decoding 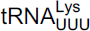 and AAG-decoding 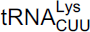 into the mammalian termination complexes does not immediately suggest how these tRNAs could differentially influence eRF1 action (**Supplementary Fig. 7D, E**). In fact, the conformation of the tRNA^Val^ in the termination complex appears to be very similar to that of tRNA^Lys^ and tRNA^Arg^ observed in stalled ribosomal complexes (**Supplementary Fig. 7F**). However, such analysis does not consider differences in the conformation of the tRNAs, nor their dynamics. Although the nature of the stop codon does not play a role in the stalling observed in the absence of New1, we note that increased readthrough is observed at weak stop codon contexts, such as UGAC, when the specific isoacceptors are present in the P site. We suggest that this arises because specific P-site isoacceptors prevent eRF1 functioning either by interfering with stable eRF1 binding and decoding and/or peptidyl-tRNA hydrolysis, and as a consequence, the stop codon can be misread by a tRNA suppressor, leading to stop codon readthrough. The peripheral binding site observed for New1 on the ribosome suggest that it does not play a direct role in promoting termination at specific tRNA isoacceptors. Instead, we propose that New1 facilitates this reaction indirectly, by promoting a conformation of the ribosome that is optimal for translation termination. Since New1 binds predominantly to the head of the 40S subunit and binds stably to non-rotated state complexes, we speculate that the ribosome may adopt non-productive conformations when specific tRNA isoacceptors are present in the P site and that by stabilizing non-rotated ribosomes with non-swivelled heads, New1 may facilitate binding of eRF1 and peptidyl-tRNA hydrolysis during translation termination.

## DATA AVAILABILITY

5PSeq sequencing data are available in NCBI Gene Expression Omnibus (GEO) repository under GEO accession GSE229473. Cryo-EM maps have been deposited in the Electron Microscopy Data Bank (EMDB) with the following accession codes: State 1 -New1-80S ribosome with A-tRNA, P-tRNA, eIF5a (EMD-19917); State 2 - New1-80S ribosome with A-tRNA, P-E*-tRNA (EMD-19918); State 3 - New1-80S ribosome with A-P-tRNA, P-E-tRNA (EMD-19925); State 4 - New1-80S ribosome with P-tRNA, E-tRNA (EMD-19923); State 5 - New1-80S ribosome with A-tRNA, P-tRNA, E-tRNA (EMD-19921); State 6 - New1-80S ribosome with A-tRNA, P-tRNA, E-tRNA (EMD-19922); State 7 - New1-80S ribosome with eRF1, P-tRNA, E-tRNA (EMD-19924).

## ACKNOWLEDGMENTS

We are grateful to Anders Byström for providing the pAG25 plasmid and Mikael Lindberg at the Protein Expertise Platform (PEP) facility at Umeå University for the construction of plasmid VHp893.

---

**Supplementary Figure 1.**
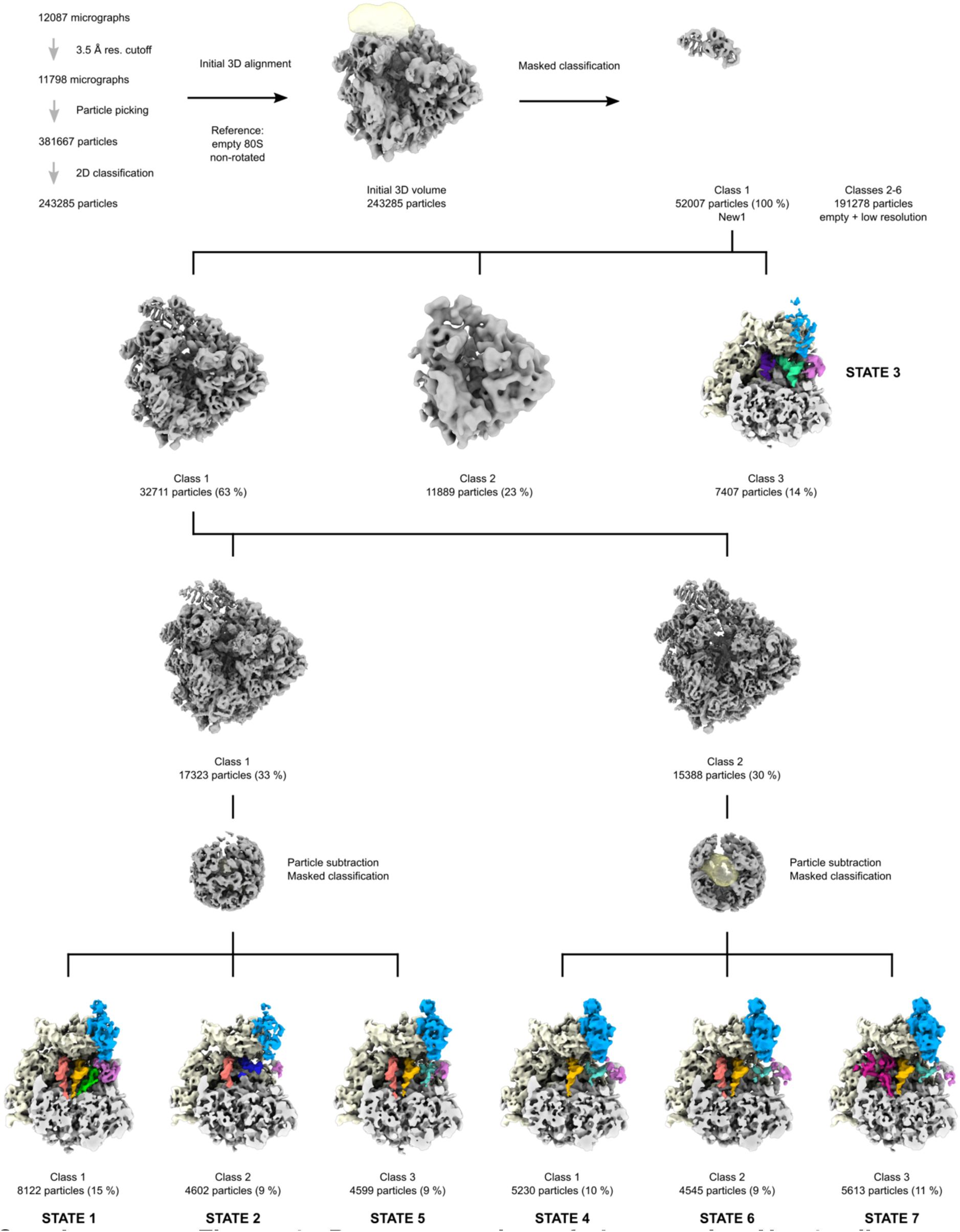
Data processing of the *ex vivo* New1p-ribosome complexes. (A) From 12,087 micrographs, 243,285 ribosomal particles were obtained after 2D classification, yielding (B) an initial 3D volume, that was (C) subjected to masked classification (mask over the New1p area). Classes 2-6 were discarded due to the presence of either vacant ribosomes or low resolution. (D) Class 1 was further 3D classified yielding State 3, and (E-F) a class (Class 1) that was further 3D classified using particle subtraction to eventually yielded States 1, 2, 4-7.

**Supplementary Figure 2.**
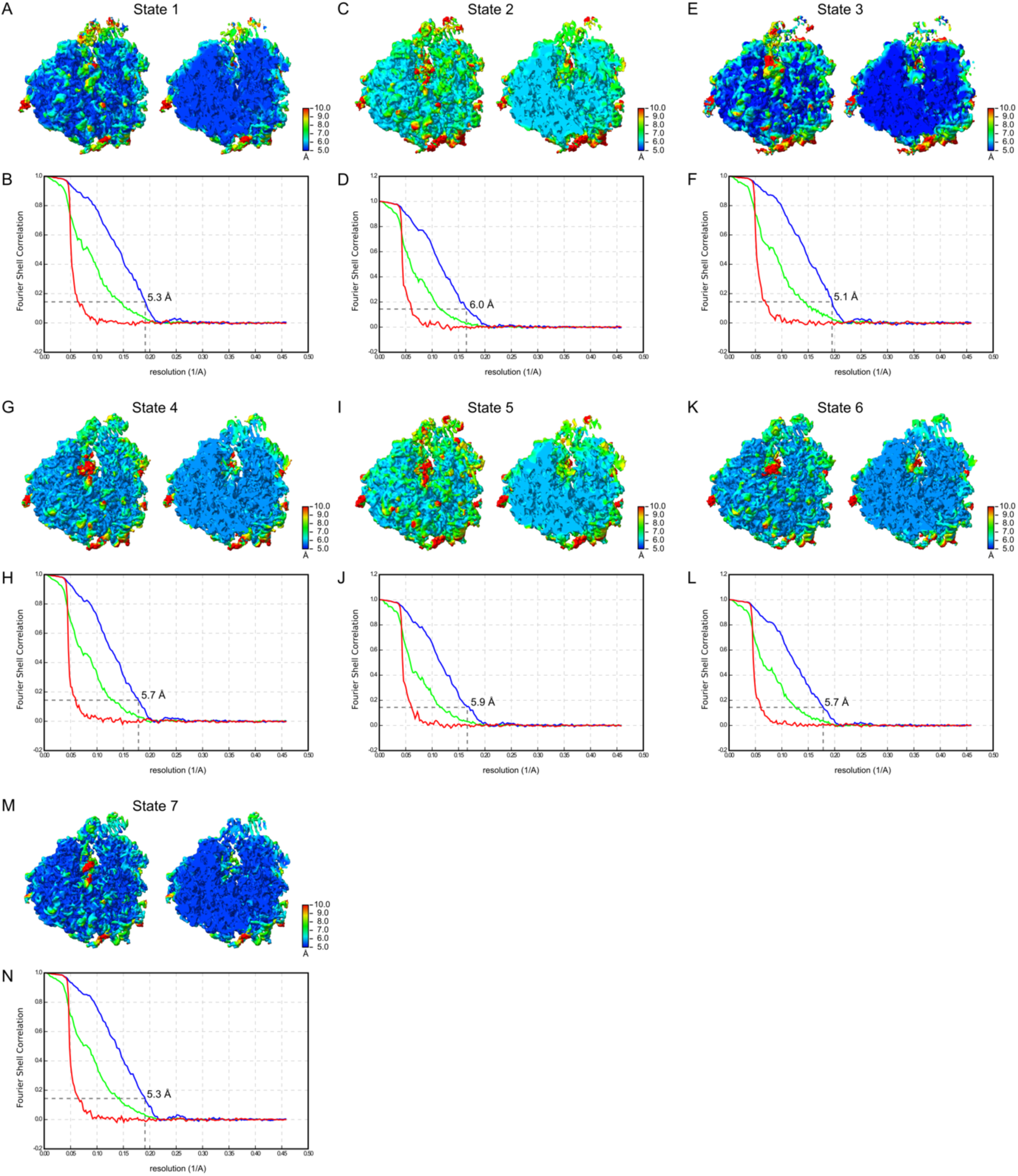
Local resolution and Fourier Shell Correlations of States 1-7. (A-M). Overview (left) and transverse section (right) of the cryo-EM map colored by local resolution is shown with the Fourier Shell Correlation (FSC) curve for (A-B) State 1, (C-D) State 2, (E-F) State 3, (G-H) State 4, (I-J) State 5, (K-L) State 6, and (M-N) State 7. The resolution scale for the local resolution is indicated for each state, and the dashed line in the FSC graph at 0.143 indicates the average resolution for each state. The different curves in the FSC graph include the masked map (green), unmasked map (blue), the phase-randomized masked map (red).

**Supplementary Figure 3.**
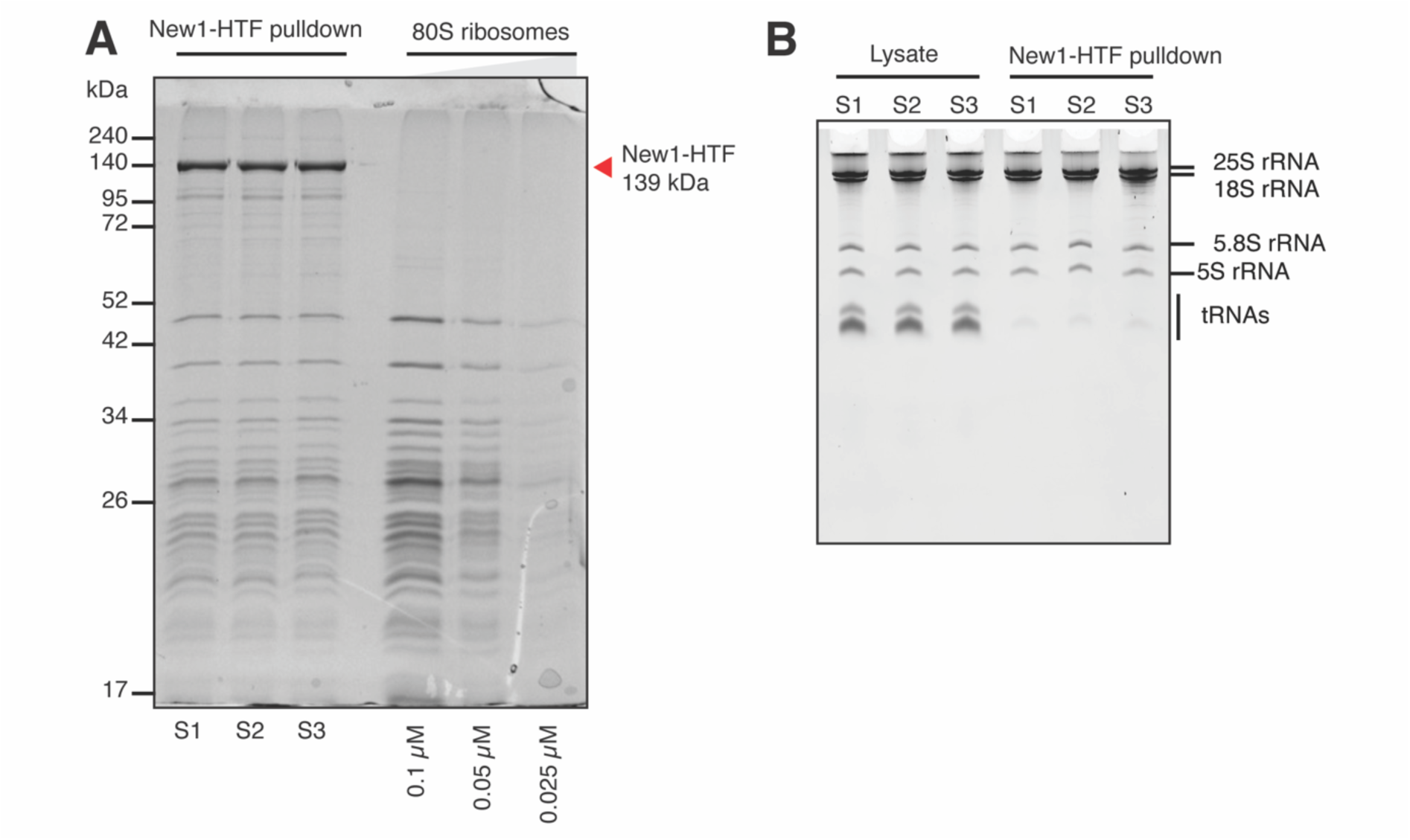
Affinity purification of New1-HTF in complex with ribosomes and associated mRNA. (A) Affinity purified samples, in triplicate, from *new1Δ* (VHY68) cells ectopically expressing New1-HTF from VHp911. The cells were grown at 20°C in SC-ura medium and the expression of New1-HTF was induced by supplementing the medium with 2 µM β-estradiol. To identify co-purified ribosomal proteins and to assay ribosomal concentration within the samples, a 2-fold serial dilution of purified *S. cerevisiae* 80S ribosomes were loaded on the gel. The samples were resolved on a 15 % SDS page gel. (B) Total RNA was extracted from the input cell lysate and the New1-HTF affinity-purified samples and resolved on a denaturing 8M urea-TBE-acrylamide gel.

**Supplementary Figure 4.**
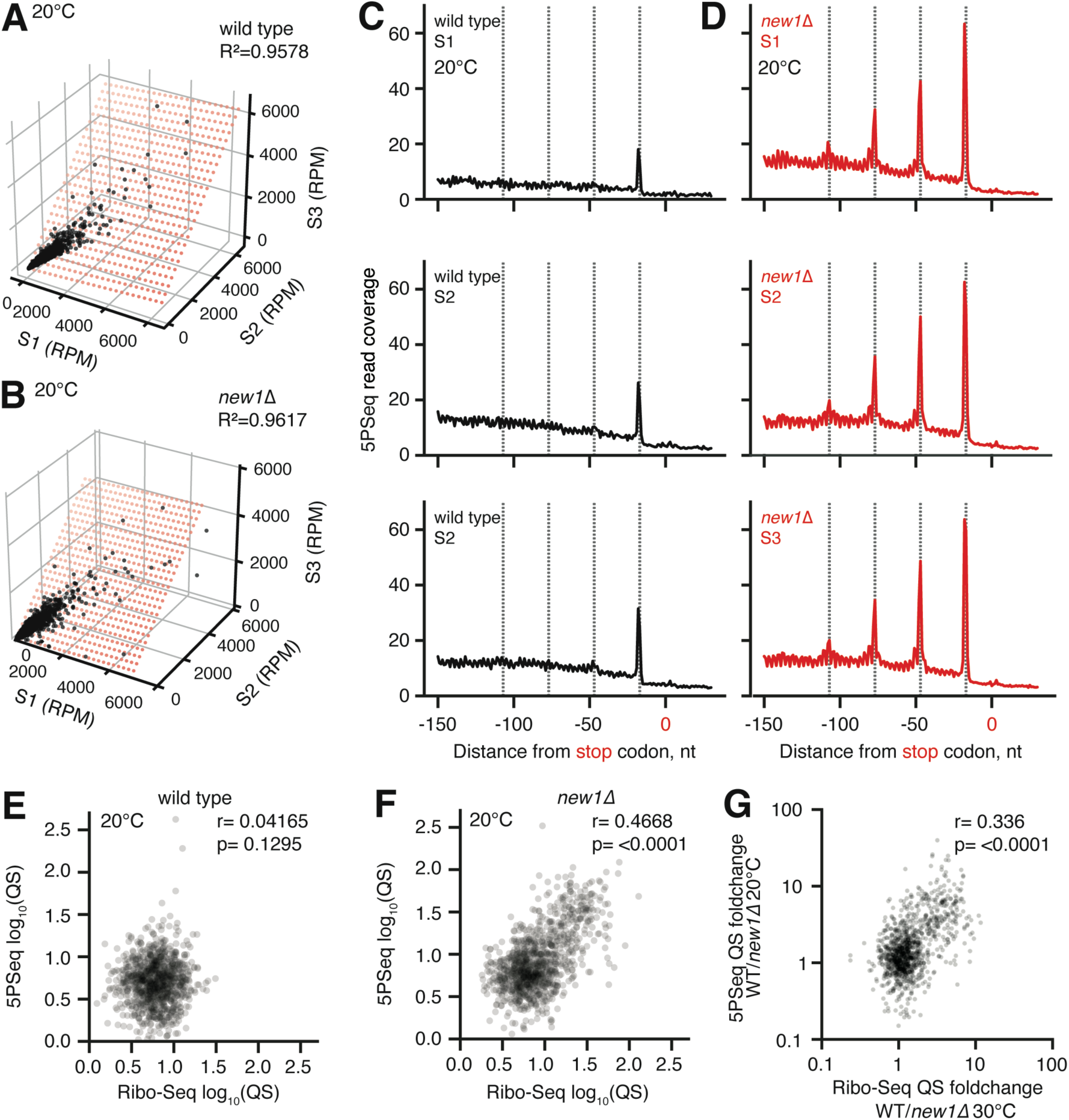
5PSeq data are highly reproducible and are in general agreement with the Ribo-Seq data from Kasari *et al*. 2019 (Kasari *et al*., 2019b), related to Figure 3. (A,B) Individual ORF read counts (RPM) for individual ORFs for three biological replicates of 5PSeq libraries constructed from wild-type MYJ1171 (A) and *new1Δ* MYJ1173 (B) strains. (C,D) Metagene analysis of three biological replicas 5PSeq libraries in wild-type (C) and *new1Δ* (D) strains (20°C). (E) Comparison of queuing scores (QS) for individual ORFs for pooled 5PSeq (this work) vs Ribo-Seq (Kasari *et al*., 2019b); both wild-type strain. (F) Comparison of pooled 5PSeq and Ribo-Seq derived queuing scores in the wild-type strain. (G) QS fold change between wild-type and *new1Δ* strains calculated for 5PSeq and Ribo-Seq datasets. Both 5PSeq and Ribo-Seq analyses were performed on datasets from three biological replicates collected at 20°C.

**Supplementary Figure 5.**
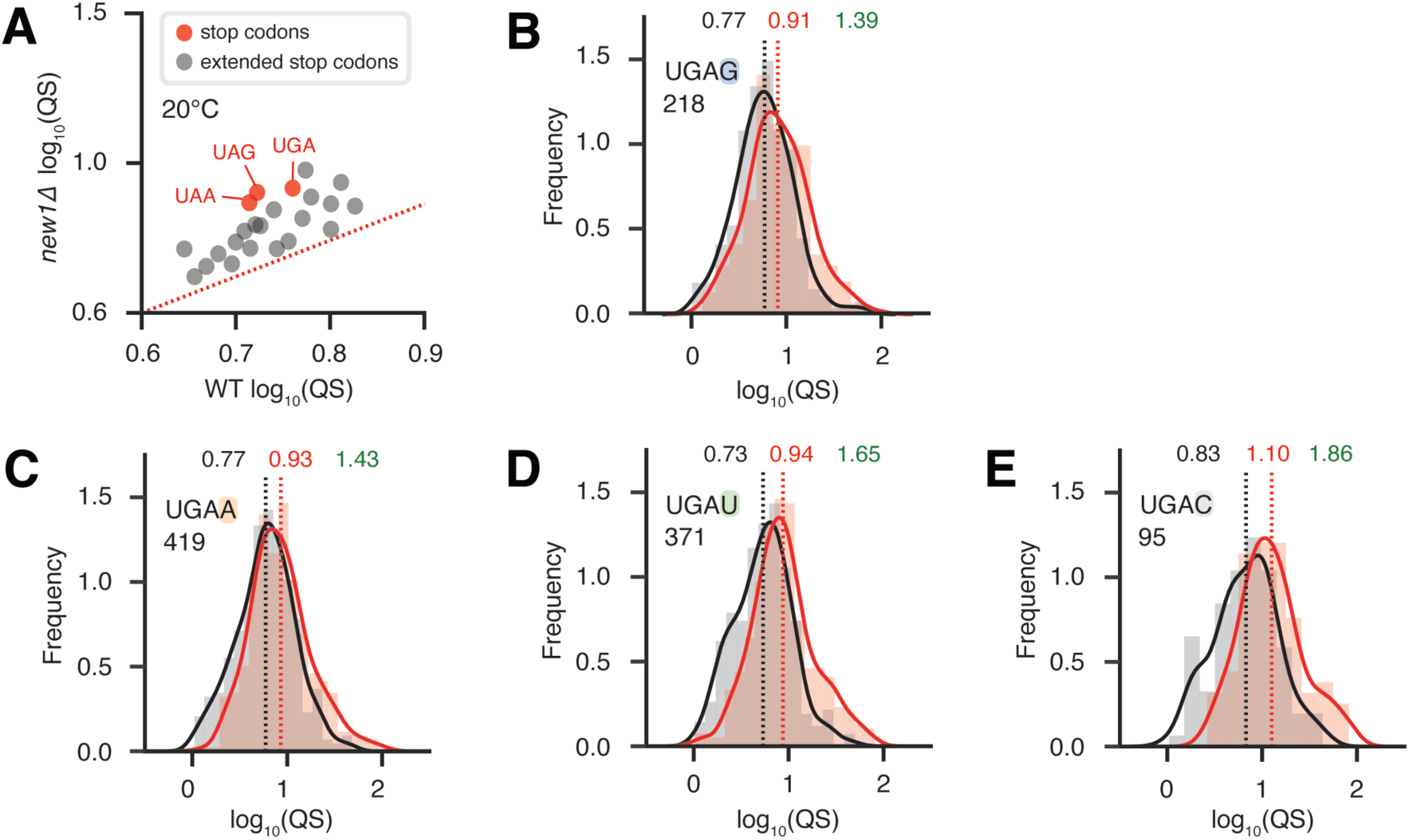
New1-dependant ribosomal queuing at the stop codon is not determined by the nature of stop codon 3’ context, related to Figure 4. (A) Mean ribosomal queuing score (QS) of ORFs in both the wild-type and *new1Δ* strains, with ORFs parsed by the nature of either extended stop codon (stop codon +1 nt), or just stop codon (grey and red circles, respectively). The red dashed line is a guide for the expected position of data points if there would be no systematic change between in QS between wild type and *new1Δ*. (**B**-**E**) Ribosome queuing score distributions sorted by the nature of the 3’ context of the UGA stop codon: (B) UAG G, (C) UAG A, (D) UAG U and (E) UAG C. Geomean QS values in the wild-type and *new1Δ* strains are given in black and red, respectively, and the QS fold change is in green. All analyses were performed on pooled 5PSeq dataset from three replicates collected at 20°C.

**Supplementary Figure 6.**
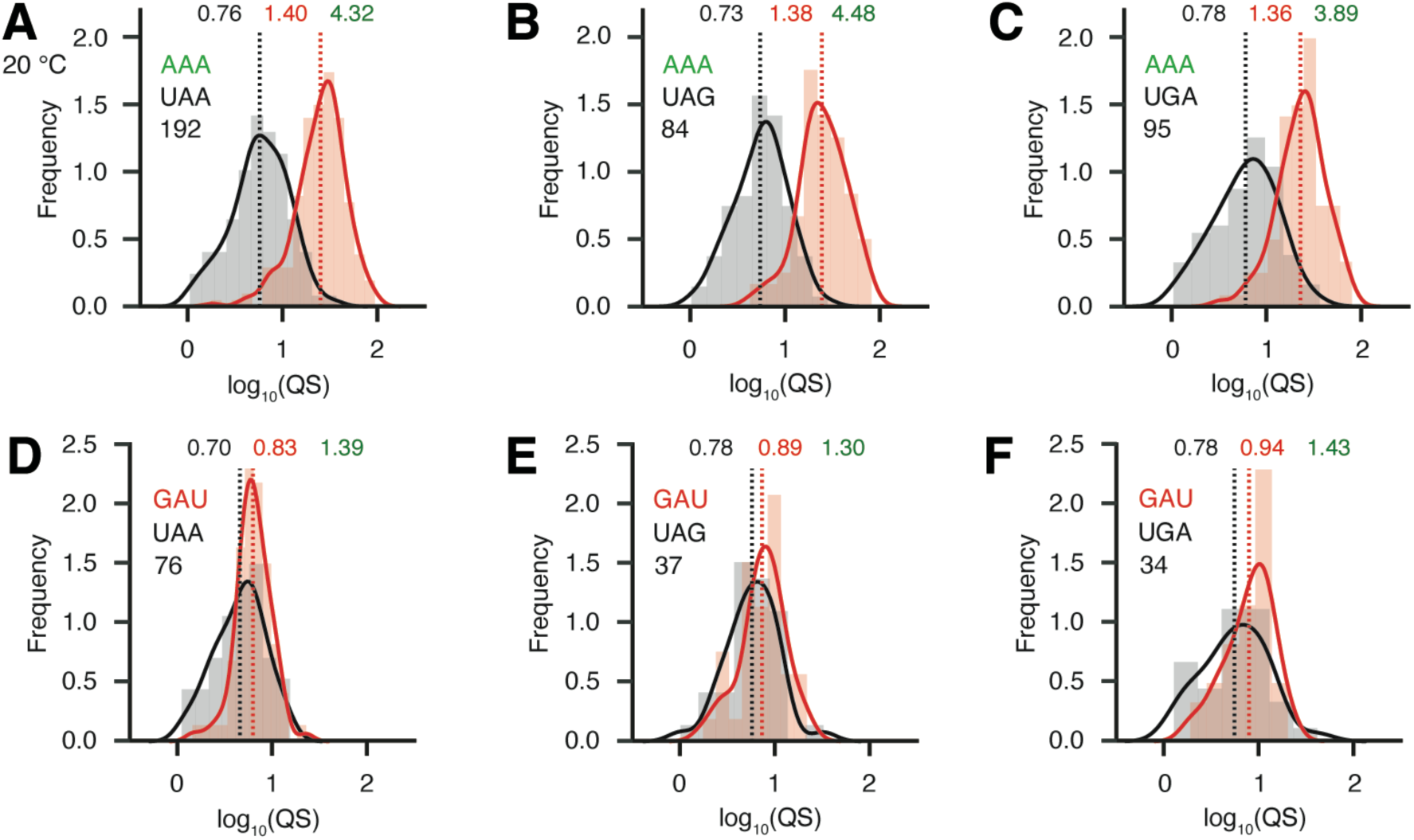
Loss of New1 results in ribosomal pile up at C-terminal AAA lysine codons regardless the nature of the stop codon, related to **Figure 4**. Ribosome queuing score distributions focussing on ORFs parsed by the nature of the C-terminal amino acid [(A-C) lysine AAA and (D-F) asparagine GAU codon] as well as the stop codon. The nature of the +4 reside was not taken into account while parsing the data. Number of included ORFs in each analysis is shown on the insert, e.g. the analysis presented on (A) was performed using 192 instances of ORFs encoding UAA preceded by AAA Lys codon. Geomean QS values in the wild-type and *new1Δ* strains are shown in black and red, respectively, and fold change in QS in green. All analysis was performed on pooled 5PSeq dataset from three replicates collected at 20°C.

**Supplementary Figure 7.**
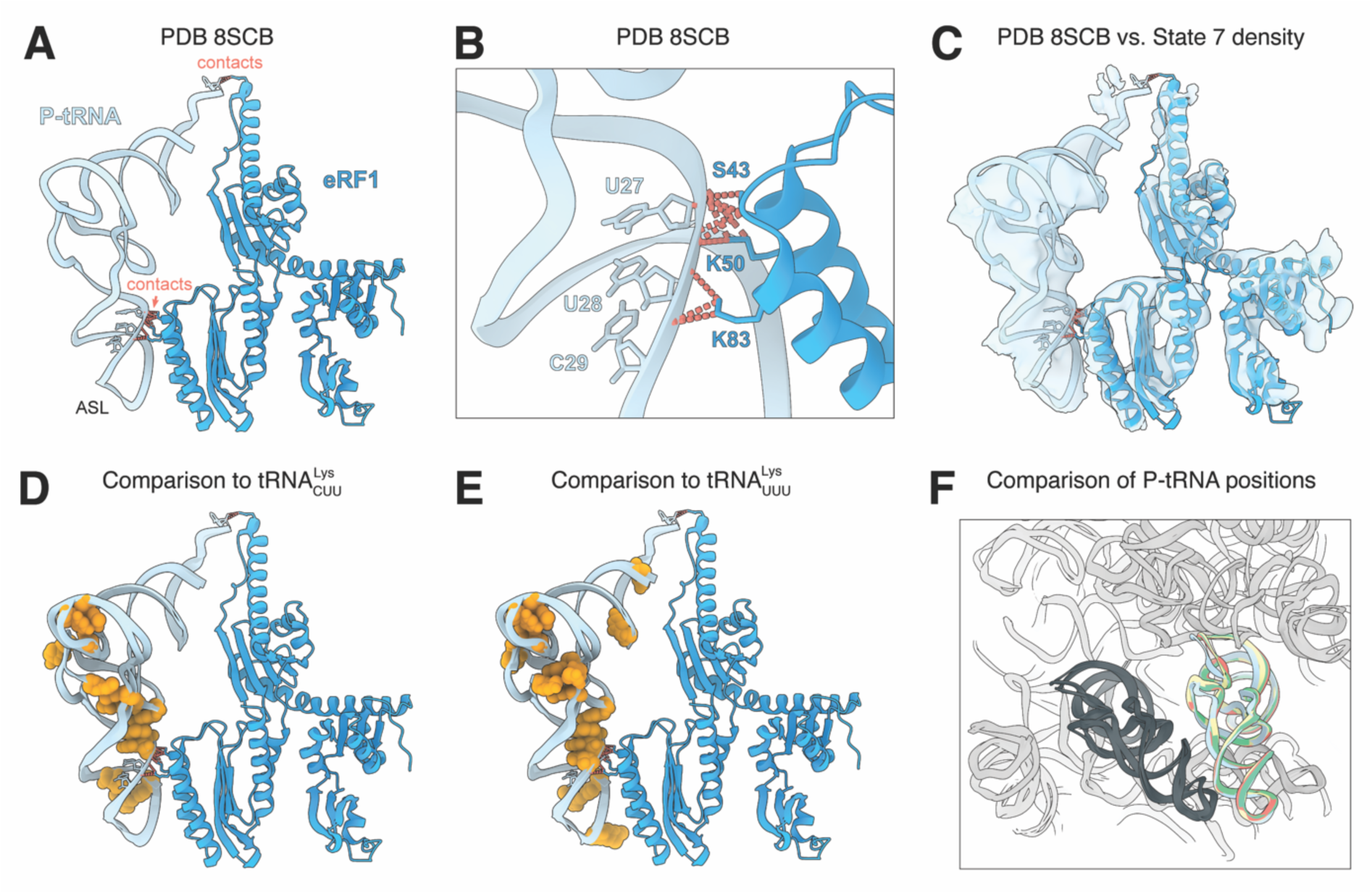
Structures of termination complexes. (**A,B**) Interaction between P-tRNA (cyan) and eRF1 (dark blue) within a mammalian (rabbit) termination complex (PDB ID 8SCB, (Coelho *et al*., 2024)) with sites of contact indicated by red dashed lines. (**C**) Alignment of P-tRNA and eRF1 from (A) into the cryo-EM density (transparent blue) of State 7 of the New1-termination complex determined here. (**D**,**E**) threading of the sequence and modifications of (D) AAG-decoding 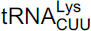 and (E) AAA-decoding 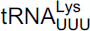 onto the P-tRNA (cyan) of the mammalian (rabbit) termination complex (PDB ID 8SCB, (Coelho *et al*., 2024)). Orange spheres indicate modifications in the respective tRNAs. (**F**) Alignment of the tRNA^Lys^ [PDB ID 6T83 (Tesina *et al*., 2020) (yellow) and 6SGC (Chandrasekaran *et al*., 2019) (cyan)] and tRNA^Arg^ (PDB ID 6T4Q (Tesina *et al*., 2020) (green) and 6T7I (Tesina *et al*., 2020) (blue)) onto the mammalian (rabbit) termination complex (PDB ID 8SCB, (Coelho *et al*., 2024)).

